# Rapidly evolving protointrons in *Saccharomyces* genomes revealed by a hungry spliceosome

**DOI:** 10.1101/515197

**Authors:** Jason Talkish, Haller Igel, Rhonda J. Perriman, Lily Shiue, Sol Katzman, Elizabeth M. Munding, Robert Shelansky, John Paul Donohue, Manuel Ares

**Affiliations:** Center for Molecular Biology of RNA, University of California, Santa Cruz, Santa Cruz, CA 95064; Department of Molecular, Cell & Developmental Biology, University of California, Santa Cruz, Santa Cruz, CA 95064; Department of Biomolecular Engineering, University of California, Santa Cruz, Santa Cruz, CA 95064

## Abstract

Introns are a prevalent feature of eukaryotic genomes, yet their origins and contributions to genome function and evolution remain mysterious. In budding yeast, repression of the highly transcribed intron-containing ribosomal protein genes (RPGs) globally increases splicing of non-RPG transcripts through reduced competition for the spliceosome. We show that under these “hungry spliceosome” conditions, splicing occurs at more than 150 previously unannotated locations we call protointrons that do not overlap known introns. Protointrons use a less constrained set of splice sites and branchpoints than standard introns, including in one case AT-AC in place of GT-AG. Protointrons are not conserved in all closely related species, suggesting that most are not under selection. Some are found in non-coding RNAs (e. g. CUTs and SUTs), where they may contribute to the creation of new genes. Others are found across boundaries between noncoding and coding sequences, or within coding sequences, where they offer pathways to the creation of new protein variants, or new regulatory controls for existing genes. We define protointrons as (1) nonconserved intron-like sequences that are (2) infrequently spliced, and importantly (3) are not currently understood to contribute to gene expression or regulation in the way that standard introns function. A very few protointrons in *S. cerevisiae* challenge this classification by their increased splicing frequency and potential function, consistent with the proposed evolutionary process of “intronization”, whereby new standard introns are created. This snapshot of intron evolution highlights the important role of the spliceosome in the expansion of transcribed genomic sequence space, providing a pathway for the rare events that may lead to the birth of new eukaryotic genes and the refinement of existing gene function.

**Author Summary:** The protein coding information in eukaryotic genes is broken by intervening sequences called introns that are removed from RNA during transcription by a large protein-RNA complex called the spliceosome. Where introns come from and how the spliceosome contributes to genome evolution are open questions. In this study, we find more than 150 new places in the yeast genome that are recognized by the spliceosome and spliced out as introns. Since they appear to have arisen very recently in evolution by sequence drift and do not appear to contribute to gene expression or its regulation, we call these protointrons. Protointrons are found in both protein-coding and non-coding RNAs and are not efficiently removed by the splicing machinery. Although most protointrons are not conserved, a few are spliced more efficiently, and are located where they might begin to play functional roles in gene expression, as predicted by the proposed process of intronization. The challenge now is to understand how spontaneously appearing splicing events like protointrons might contribute to the creation of new genes, new genetic controls, and new protein isoforms as genomes evolve.

## Introduction

Eukaryotic genes are often split by intervening sequences called introns that are removed during and after transcription by the spliceosome and associated splicing proteins. Although much is known about the biochemical mechanisms of intron recognition and splicing [1-3], a clear understanding of the events and processes that explain the appearance and persistence of introns during the evolution of eukaryotic genomes remains elusive [4, 5].

As a necessary step in the expression of most extant eukaryotic genes, splicing has been exploited by evolution in at least two main ways. One allows diversification of the structure and function of the RNA and protein products of a gene by producing multiple distinct mRNAs through alternative splicing [2]. A second allows changes in gene expression through nonsense-mediated decay (NMD), whereby alternative splicing can lead to either functional mRNA, or to transcripts with premature stop codons that are degraded, providing developmental on-off control, or stable homeostatic expression settings [2]. The complex gene architecture of multicellular organisms, as contrasted with the simpler gene architecture in many single-celled eukaryotes, has prompted widespread speculation that alternative splicing is responsible for emergent complexity in metazoans. Although it contributes in complex and critical ways to gene function and regulation in extant eukaryotes, how splicing came to reside so pervasively in the eukaryotic lineage remains to be explained [4-6].

Until recently, gain or loss of introns has been detected by comparing closely related genomes. Many such “presence/absence” variations are inferred to be intron loss, in which reverse transcription of a spliced RNA, followed by homologous recombination of the intronless cDNA back into the gene of origin, erases the intron [6, 7]. Several mechanisms for the gain of new introns have been proposed (for review see [8, 9]). For example, single nucleotide changes that create new splice sites (and thus new introns) can lead to “intron sliding” or new alternative splicing events [10]. “Exonization” of an Alu sequence in a large intron can lead to inclusion of a new exon and splitting of an intron into two smaller introns [11]. These intron gain mechanisms rely on pre-existing local splicing events, and represent intron diversification, rather than de novo intron creation at sites where no intron previously existed.

De novo intron creation appears to occur by two main pathways, “intron transposition” whereby an intron at one location is copied and inserted at a new location, and “intronization”. First described in the marine alga *Micromonas* [12], transposition of introns called “Introner Elements” appears to have expanded an intron repeat family in some lineages [13-17] perhaps through an “armed spliceosome” carrying an intron-lariat RNA which is then reverse spliced into an mRNA (which then must be converted to cDNA to return to the genome at a new location, [6, 9]). More recently, intron transposition through an RNA intermediate has been documented in *S. cerevisiae*, supporting the idea that reverse splicing may operate to spread introns [18]. Other models suggest that introns transposition may arise by the action of DNA damage repair [19] or non-autonomous DNA transposons [20].

A distinct pathway for *de novo* intron creation is called “intronization” whereby mutations arise either through drift [4, 8, 21], or other sequence changes [22] to create sequences recognized as introns by the splicing machinery. As a genome sequence distant from other introns drifts, it may accumulate mutations that by chance allow its transcripts to be recognized by the splicing machinery and spliced. This process is thought to occur gradually over evolutionary time, generating sequences that exhibit properties of both exons and introns that are often alternatively spliced through several, different, weak splicing signals [4, 23]. Whether these sequences evolve to become bona fide introns depends on whether their removal through splicing provides a fitness advantage.

Splicing in the *S. cerevisiae* genome appears to have been streamlined by evolution, such that about 5% of genes have introns, and most that do have only one. Despite their scarcity in genes, introns appear in about 25% of transcripts when cells are growing in rich medium [24, 25]. More than a third of annotated introns are found in genes for ribosome biogenesis (ribosomal protein genes, RPGs), and their mRNAs account for 90% of the splicing performed in rapidly growing cells [25, 26]. This unusual distribution of introns in a highly expressed class of genes with shared function presents both challenges and opportunities for studying integration of splicing into core cellular regulation. For example, repressing transcription of the RPGs increases the efficiency of splicing for the majority of non-RPG introns and can suppress temperature-sensitive spliceosomal protein mutations [27, 28]. Based on this we proposed that relieving pre-mRNA competition by reducing RPG expression frees the spliceosome to process less competitive, splicing substrates it normally ignores. This phenomenon has been shown to contribute to the efficiency of meiotic splicing [28, 29], as well as regulation of Coenzyme Q_6_ synthesis [30], whereby regulated repression of RPGs potentiates splicing control mechanisms at other genes.

In this study, we use rapamycin to repress RPGs and create hungry spliceosome conditions in three related yeast species that have diverged over ~10-20 million years [31], and find 163 locations in the *S. cerevisiae* genome that are substrates of the spliceosome, but distinct from the current set of annotated introns. Other studies have also found undocumented splicing events in yeast [32-39], including alternative splicing of known introns. Here we focus strictly on splicing events not associated with a known intron, which we call protointrons. To better understand intron creation, we distinguish, standard introns from protointrons as follows. Standard introns are efficiently spliced under normal growth conditions (median efficiency ~93% spliced), highly conserved in related *Saccharomyces* species, and have known functions in gene expression (93% reside in protein coding regions). Protointrons on the other hand generally splice with very low efficiency (median efficiency ~1% spliced), are often specific to different *Saccharomyces* species, and most importantly do not have a clear role in correct gene expression. These protointrons are found in mRNAs as well as noncoding RNAs such as promoter-associated transcripts, CUTs, SUTs, XUTs and other noncoding RNAs [40-48]. Related yeasts *S. bayanus* and *S. mikatae* also have protointrons, but most are species-specific, indicating that protointrons appear and disappear during evolution. This suggests that protointrons arise initially through genetic drift and provide the raw material for intronization in order to enable intron creation through a mechanism that is distinct from gene duplication or intron transposition [4]. While only a very tiny fraction of protointrons might ever evolve into new standard introns, we observe several, more efficiently spliced protointrons that appear to be more advanced along the intronization pathway. This intermediate class of introns tend to occur in 5’UTRs where they might buffer the negative effects of RNA secondary structure or micro-ORFs (uORFs) on translation, however there is currently no evidence that these have any adaptive value. This work reveals the extent to which the spliceosome recognizes and splices intron-like sequences, thus expanding the information contained in the genome. This contribution of the spliceosome to the information content of the transcriptome may enhance the rate at which new genes and regulatory mechanisms appear in eukaryotes.

## Results

### Deeper RNA sequencing of a nonsense-mediated decay deficient strain confirms a class of rare splicing events in *Saccharomyces cerevisiae*

In experiments where we observed increases in splicing efficiency of standard introns after repression of RPG expression [28], we also observed unannotated splicing events, distinct from known introns, whose splicing efficiency improved. In addition, inventive new RNAseq methods designed to capture branchpoints [32, 34] have provided evidence for splicing events elsewhere in the transcriptome. Some of these events appear to be activated in response to stress [32], which down-regulates RPG expression and creates hungry spliceosome conditions. Still others are more readily detected during meiosis [32, 39], or when cells are deleted for RNA decay pathway components that degrade unstable transcripts [32, 33, 38]. Our interest in understanding the evolution of splicing prompted us to focus on these distinct new introns. To capture more of them, we obtained additional RNAseq data that included non-polyadenylated RNAs and RNAs sensitive to nonsense-mediated decay. We made four libraries, one each from rRNA-depleted RNA from untreated (0 min) and rapamycin treated (blocks nutrient signaling and represses RPGs, 60 min) replicate cultures of a yeast strain deficient in NMD (*upf1Δ*, [49]). We obtained more than 300 million reads that show excellent between-replicate coherence in gene expression changes (Fig. S1A). These data confirm our previous observation [28] that splicing of a majority of standard introns in non-RPGs increases after rapamycin treatment. A splicing index relating change in the ratio of junction/intron reads over time (SJ index = log2[junctions-t60/intron-t60] – log2[junctions-t0/intron-t0]) increases for most introns in transcripts whose total transcript levels change less than two-fold during the experiment (Fig 1A, blue circles, Table S1, NB: RPGs are repressed >2 fold and are excluded). In addition to the standard introns, we observe more than 600 splicing events that result from use of alternative 3’ or 5’ splice sites overlapping the standard introns (see also the study by Douglass et al. [38]). Some of these splicing events have been previously characterized [33, 36, 37], and produce out of frame mRNAs that are more easily detected in the *upf1Δ* strain due to the loss of NMD (Table S2). Because our interest here is in introns appearing at novel locations, we have not studied the splicing events that overlap the standard introns any further. At this sequencing depth, a set of splicing events that do not overlap any standard intron is evident (Table 1 and Table S3). These are supported by reads that span sequences with known characteristics of introns but occur at much lower frequencies than those for standard introns. As is the case for standard introns, splicing of the majority of these nonstandard introns increases upon rapamycin treatment (Fig 1A, orange circles, positive values indicate increased splicing), suggesting they are also in competition with RPG pre-mRNAs for the spliceosome.

**Figure 1.**
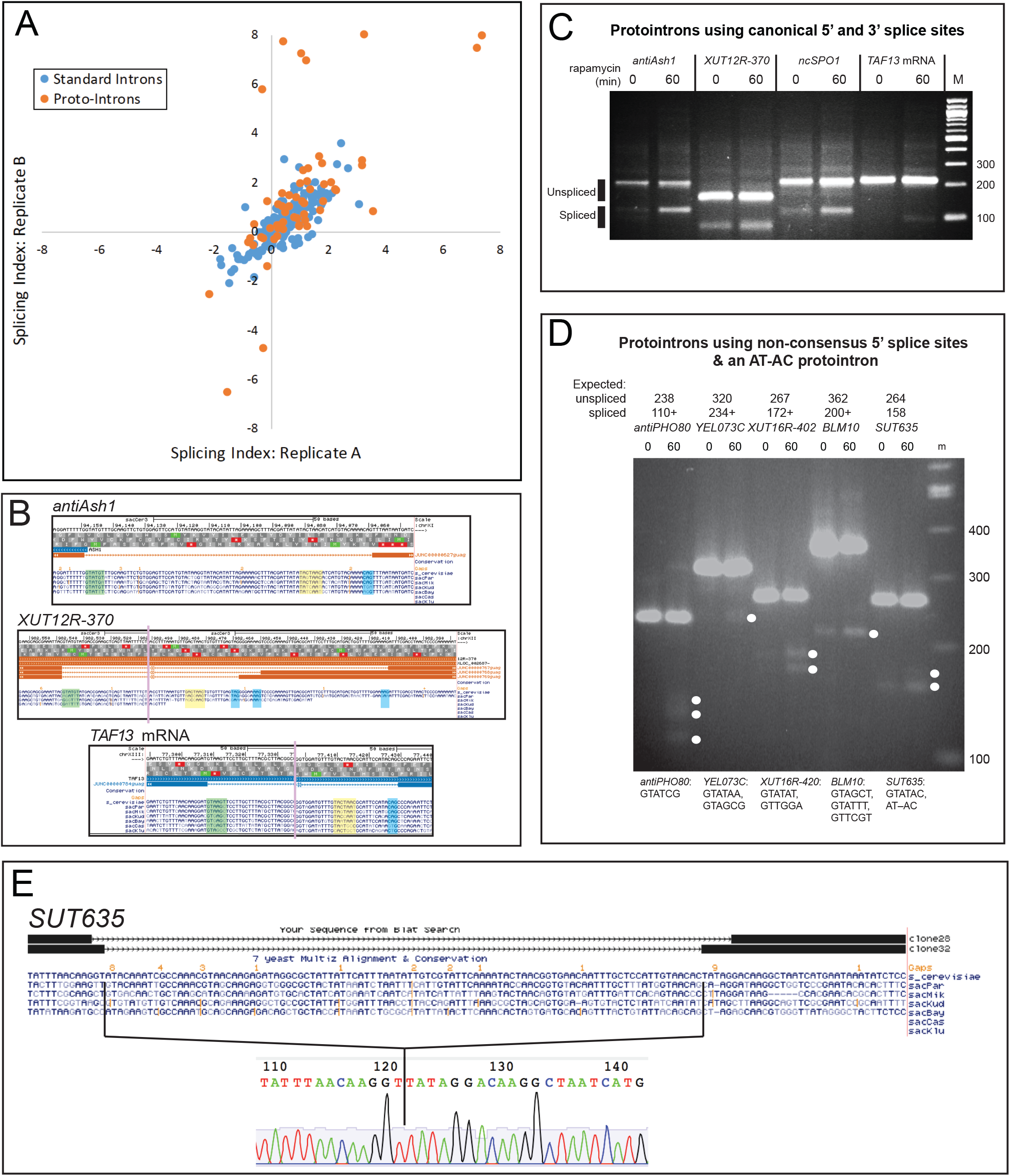
Identification and Validation of Splicing at Unannotated Genomic Locations. RNA sequencing reads corresponding to spliced RNAs defined as described in the text were used to identify and measure splicing. Reads (~300M) were obtained from untreated cells and cells treated for 60 minutes with rapamycin from two biological replicate experiments. **(A)** Splicing efficiency of many introns improves after rapamycin treatment as judged by the log2 fold change in the ratio of splice junction reads to intron reads (splicing index; see Methods) for replicate experiments. Standard introns with ≥35 total junction reads are shown as blue dots. Unannotated and non-overlapping splicing events (protointrons) with ≥50 total junction reads are shown as orange dots. Data points in quadrant I indicate introns in which splicing improved after treatment with rapamycin in both replicate experiments. **(B)** Genomic alignment and lack of conservation for three example protointrons. Protointrons in a divergent upstream transcript *antiASH1*, a XUT *XUT12R-370*, and an mRNA for *TAF13* are shown. The 5’ ss is green, the branchpoint sequence is yellow, and the 3’ ss is blue. The vertical bars indicate where additional intron sequences are not shown. **(C)** RT-PCR validation of protointron splicing and increased splicing after rapamycin is shown for the protointrons in (B), and for a non-coding vegetative cell transcript of a meiotic gene *ncSPO1*. PCR products corresponding in size to spliced (predicted based on RNAseq read structure) and unspliced RNAs are labeled. Splice junctions were confirmed by sequencing cloned PCR products. **(D)** Validation of protointron splicing through 5’ ss not observed in standard introns. Junctions were validated by sequencing cloned PCR products. **(E)** A protointron in *SUT635* uses both GT-AG and AT-AC splice sites. The sequences of two cloned PCR products from *SUT635* are aligned to the genome (above) and the sequencing trace from the clone representing the use of AT-AC junctions is shown (below).

**Table 1.**
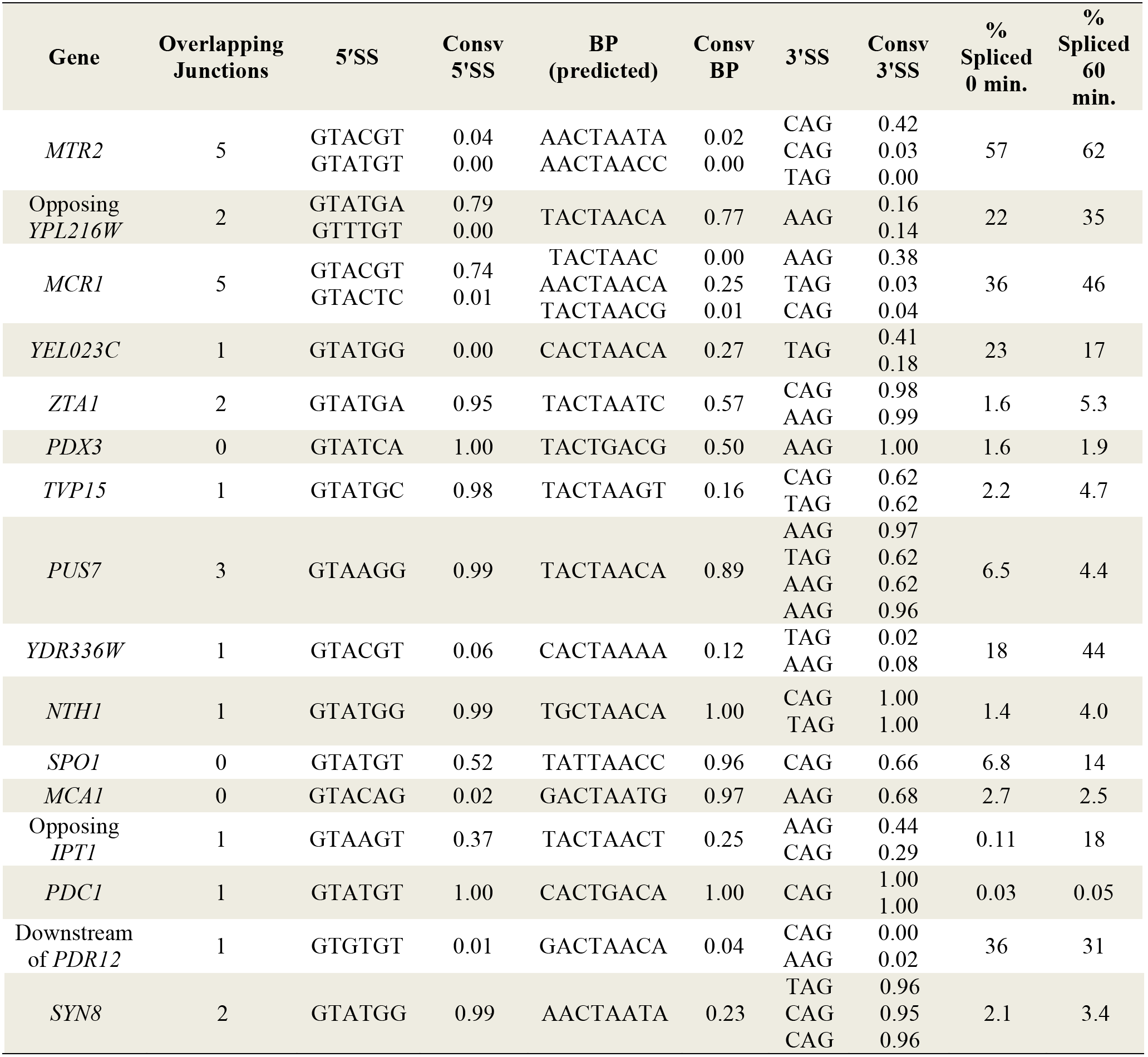
Easily detectable protointrons in *Saccharomyces cerevisiae upf1Δ* mutant cells

### Identification and validation of protointrons

To extract and validate nonstandard splicing locations from the RNAseq data, we inspected reads that span and are missing genomic sequences bounded by GT or GC on the 5’ side and AG on the 3’ side, aggressively filtering out those that were unlikely to have been generated by the spliceosome (Table S4, see also Methods). For example, we ignored reads that are abundant in only one library due to spurious PCR-derived “jackpot” amplification, or that are incorrectly mapped as spliced over naturally repeated sequences. We also filtered out reads that appeared fewer than three times, spanned sequences with no discernable match to a relaxed and appropriately positioned branchpoint consensus sequence (RYURAY, >45 from a 5’ss, >7 from a 3’ss), or that use a GAG 3’ ss (although some of these may be true). Finally, in order to focus on new intron locations, we separated out reads that overlap known standard introns (these are found in Table S2). Finally, we merged overlapping alternative splice sites observed at each new intron location into one. From this sequencing experiment, we identify 226 splicing events at 163 intron locations in the *Saccharomyces cerevisiae* genome that do not overlap with standard introns. We call these protointrons (Table 1 and Table S3, NB: we have reclassified some previously annotated introns as protointrons based on their less efficient splicing and lack of conservation in close relatives, see below).

To characterize protointrons as a distinct class of splicing events, we inspected the alignments of many individual protointrons (for example Fig 1B) and validated more than a dozen of them by RT-PCR, cloning, and Sanger sequencing (Fig 1B and C, Table S5). These events map to a diversity of transcribed locations, including mRNAs, a variety of non-coding RNAs (CUTs, SUTs, XUTs, etc.), and within telomeric Y’ repeats. 39% of protointrons can be found entirely within a coding region (e. g. *TAF13* Fig 1B, C, Fig S1B and E), whereas others reside in noncoding regions or within RNA antisense to a standard gene (29%, e. g. *antiASH1*, Fig 1B, C, Fig S1B and C). Often alternative splice sites are observed, for example in the noncoding RNA *XUT12R-370* antisense to *TUS1* (Fig 1B, note only the product derived from use of the downstream 3’ ss is visible on the gel in Fig 1C, Fig S1D). An internally initiating RNA from the meiotic *SPO1* gene expressed only during vegetative growth has a protointron (Fig 1C, Fig S1F). Protointrons can also be found crossing boundaries from the coding region to either UTR in mRNAs (17% in 5’UTR^coding; 1% in coding^3’UTR), and still others can be found completely within a UTR (9% in 5’UTR; 4% in 3’UTR). In many cases, excision, cloning and sequencing the faint band near the size predicted by the RNAseq reads identifies additional alternatively spliced forms not observed by RNAseq. For the four events shown in Fig 1C, we successfully confirmed splice junctions indicated by RNAseq (Table S3).

Several of the protointrons predicted by the RNAseq data appear to use unusual 5’ splice sites (5’ ss) not anticipated by examination of standard introns. Standard intron 5’ ss show strong conservation of the G residue at position 5 (G5, underlined here: GUAY<u>G</u>U), which contributes to intron recognition through interactions first with U1 snRNA and later with U6 snRNA [50, 51]. The 5’ ss consensus in mammals has a less strongly conserved G5, and a subset of mammalian introns use G at position 6 instead [52]. Validation tests of several protointrons whose 5’ ss lack G5 show that they are authentic products of splicing (Fig 1D, Fig S1G). During this effort we also detected splicing at an AT-AC junction in the noncoding RNA *SUT635* (Fig 1E). Although yeast does not have a minor (U12) spliceosome, some major (U2) spliceosomal introns use AT-AC junctions [53], and mutation of a standard yeast intron shows that AT-AC is the most efficient splice junction dinucleotide combination after GT-AG [54]. Alternative AT-AC junction use has been reported for the standard intron (a normal GT-AG intron) in *RPL30* [32], however no standard yeast intron uses AT-AC junctions normally. The appearance of an AT-AC protointron in *SUT635* suggests that AT-AC introns may represent an alternative path to evolution of standard introns.

### Distinct features of protointrons: sequence, conservation, size, and splicing efficiency

To begin contrasting the features of protointrons and standard introns, we evaluated evolutionary conservation, the most obvious difference. To analyze conservation around splicing signals of both intron classes, we extracted a sequence window surrounding the 5’ss, predicted branchpoint, and 3’ss from each standard intron and protointron and compared them. Position-specific weight matrix-based logos of the splicing signals (Fig 2A) reveal that protointrons use a more divergent collection of splicing signals than do standard introns. As noted above, G5 of the 5’ss is a prominent feature of the standard yeast intron 5’ss but is less well represented among protointron 5’ss. Similarly, the predicted branchpoint sequences of protointrons lack strong representation of bases to either side of the core CUAA of the UACUAAC consensus for standard introns. Branchpoints of standard introns are enriched for Us upstream of the consensus and for A just downstream, and neither of these context elements are prominent in the predicted branchpoints of protointrons. The 3’ss are more similar, but as with the branchpoint, the U-rich context detectable upstream of the standard intron 3’ss is not observed in protointrons. The more diverse collection of splicing signals used by protointrons is similar to that of the overlapping alternative splice sites of standard introns [33], and a mixed set of overlapping and distinct potential introns detected using a branchpoint sequencing approach [32]. Furthermore, we note that protointrons use a less constrained set of 5’ splice sites than standard introns as compared to the branchpoint and 3’ss sequences. This may reflect important interactions between the pre-mRNA cap binding complex and the U1 snRNP during earliest steps of spliceosome assembly [55-58] and suggests these robust interactions allow greater drift of the 5’ss than other splicing signals. This is also consistent with our observation that 26% of protointrons overlap 5’ UTRs as compared with 5% for 3’ UTRs and agrees with a prediction of the intronization model developed for metazoans [8, 59]. We conclude that protointrons use a wider variety of branchpoints and splice sites than do standard introns in *S. cerevisiae* and hypothesize that protointrons may evolve toward standard intron status by acquiring mutations that enhance the context and match to the consensus of the core splicing signals, in part to increase their ability to compete with RPG pre-mRNAs [28].

**Figure 2.**
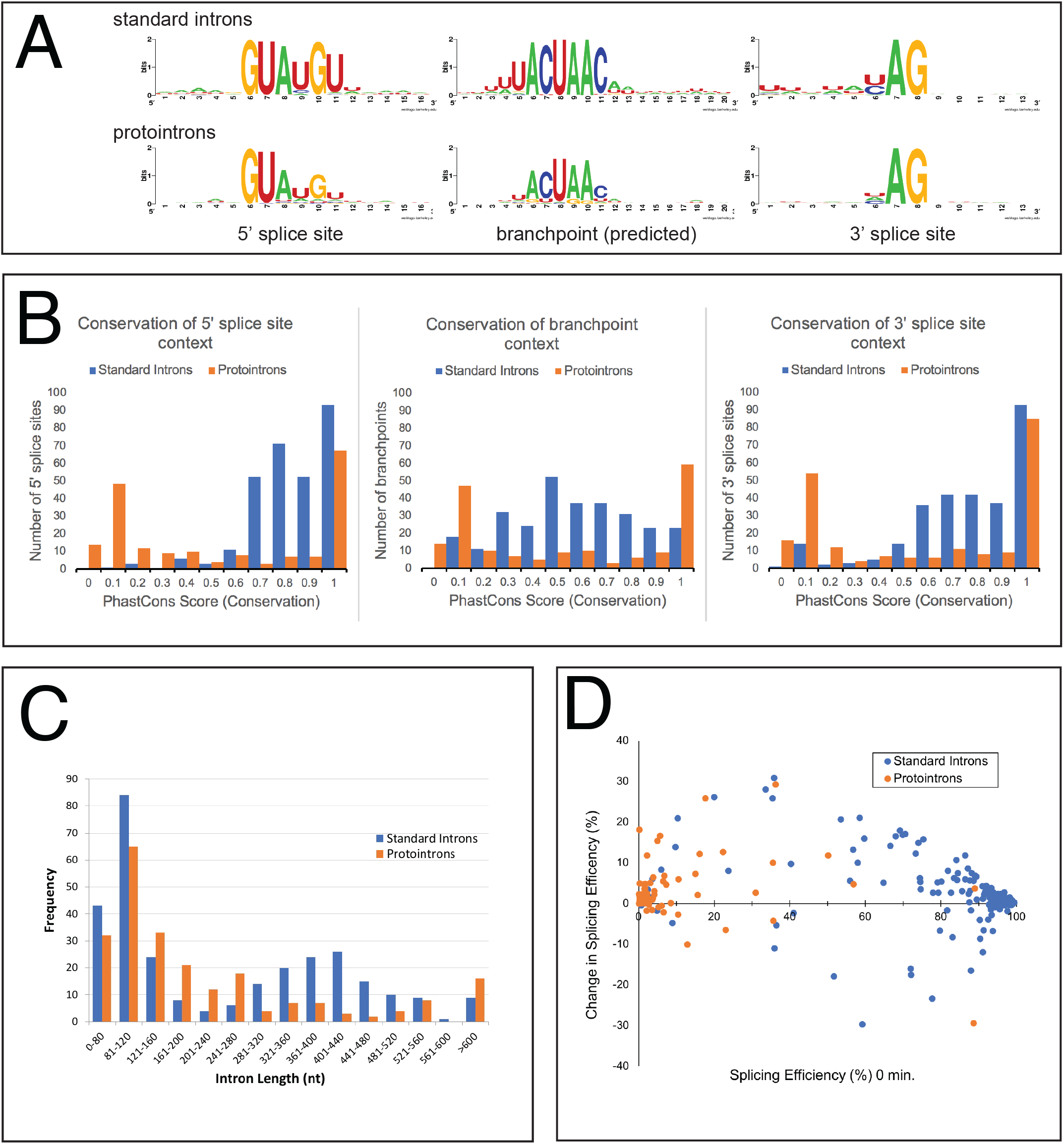
Differences between Standard Introns and Protointrons. **(A)** Splice site and branchpoint sequences of protointrons are less constrained in sequence than the standard introns. Weblogos representing position-specific weight matrices of the 5’ ss, branch points, and 3’ ss of the standard introns (top) and the protointrons (bottom) are shown. **(B)** The pattern of sequence conservation in the context of protointron splicing signals (orange bars in each panel) is bimodal as compared to the standard introns (blue bars in each panel). Histograms of the standard introns (blue bars) and the protointrons (orange bars) showing the distributions of PhastCons scores in windows containing the 5’ ss, branchpoints, and 3’ ss of the standard introns and protointrons. See text. **(C)** The size distribution of protointrons is distinct from that of the standard introns. Histograms show the distribution of intron sizes for the standard introns (blue bars) and the protointrons (orange bars). A Kolmogorov-Smirnoff test indicates the two distributions are different (*D* = 0.22, p value ≤ 10^-4^). **(D)** Protointrons are much less efficiently spliced than standard introns. The scatter plot shows the relationship between splicing efficiency in untreated (time 0) cells and the change in splicing efficiency after one hour in rapamycin. Standard introns are shown in blue, protointrons in orange.

With the exception of the protointrons found entirely within protein coding sequences, typically at least one (and often all three) of the splicing signals for a given protointron in *S. cerevisiae* is imbedded in sequence that is not conserved in closely related *Saccharomyces* species (Fig 1B, E, Fig S1C, F). To analyze whether this is a distinguishing characteristic of protointrons, we recorded the average phastCons score (range between 0, evolving as not conserved, and 1, evolving as highly conserved, [60]) of the nucleotides within sequence windows containing the 5’ss, the branchpoint, and the 3’ss of the standard introns and protointrons and plotted them (blue bars, standard introns; orange bars, protointrons, Fig 2B). Many standard introns are embedded in conserved protein coding sequences, and thus the splice sites and their immediate exon context are also conserved, such that the average phastCons scores for both the 5’ and 3’ss windows rise above 0.7 (Fig 2B). The branchpoints of standard introns are also conserved but have a broader and lower score distribution, because constraining protein coding exon sequences are not usually found near the branchpoints of standard introns. In contrast, the distributions of phastCons scores for each of the protointron splicing signals is clearly bimodal, meaning that protointron splicing signals are either highly conserved, falling within protein coding sequence, or are poorly conserved, falling in UTRs or intergenic regions (Fig 2B). This distribution illuminates the sequence landscape within which most protointrons arise. Since the strongly transcribed regions of the genome code mostly for protein, and transcription is a prerequisite for splicing, protointrons tend to appear in or span less well conserved noncoding RNA sequences such as UTRs and ncRNAs (Supplemental Figure S1B). This distinguishes protointrons from standard introns and suggests that most of the protointrons we detect have appeared only recently in the *S. cerevisiae* genome, when transcribed non-protein coding regions acquire intron-like features by mutation.

Standard introns show a bimodal length distribution, with peaks at about 100 nt and 400 nt ([61], Fig 2C). In contrast, the distribution of protointrons is on average shorter, with a single main peak at around 100 nt in length, and few larger than 300 nt. These distributions are significantly different (Kolmogorov-Smirnoff test, p ≤ 10^-4^), suggesting that if protointrons evolve into standard introns, they may become longer by acquiring additional sequence features that enhance their recognition by the spliceosome. Many of the larger standard introns are found in RPGs [61], where secondary structures and other long distance RNA-RNA interactions promote efficient and accurate splicing [62, 63]. The elaboration of such structures during evolution of increased splicing efficiency may explain the increased intron length that characterizes the large intron class in yeast.

Protointrons are less efficiently spliced than standard introns (Fig 2D, Tables 1, S1 and S3). The vast majority of standard introns are spliced at greater than 80% efficiency (median ~ 93%) by comparison of splice junction reads to intron base coverage. Exceptions include meiotic introns whose efficient splicing may require repression of RPGs or the expression of a meiosis-specific splicing factor like Mer1 [27-29]. In contrast, most protointrons have splicing efficiencies below 20% at best (median ~ 1%). A few protointrons, such as the introns in the *S. cerevisiae MTR2*, *USV1*, and *MCR1* genes or the *S. bayanus YTA12* gene, are uncharacteristically well spliced, suggesting that they may be transitional intron forms or species-specific standard introns (see below). Splicing improves for most protointrons and standard introns after rapamycin treatment, however some standard introns appear to show reduced splicing, suggesting splicing repression in response to rapamycin at those introns.

### Coding regions are depleted of sequences required for splicing

Protointrons that emerge within coding regions (39%, Supplemental Figure S1B) might disrupt gene expression by reducing mRNA levels or creating toxic proteins. Since protointrons emerge readily from nonconserved sequence (Fig 2B), we wondered whether the appearance of protointrons within ORFs most often reduces fitness, and thus whether the frequency of splice site and branchpoint sequences might be lower than would be expected by chance in protein coding regions. In rare cases such introns might allow advantageous mRNA regulation through NMD [5, 64] to emerge (as appears to have been the case with a recently evolved standard intron in *PRP5*, [65]) or make mRNA for beneficial alternative proteins (as may be the case for *PTC7*, [30, 66, 67]; and *MRM2* [37], see below), which have in-frame conserved coding sequences through their introns. Analysis of several diverged genomes by Farlow et al. revealed that the consensus 5’ ss sequence is significantly underrepresented in the coding strand of genes compared to the noncoding strand [68]. To test whether *S. cerevisiae* coding sequences might be depleted of splicing signals, we compared their frequency in the ORF set of the extant *S. cerevisiae* genome with that in 10,000 synthetic ORF sets derived by randomizing the order of codons for each ORF. This approach maintains ORF length, integrity, GC content, and codon usage in the permuted ORF sets while generating partially randomized nucleotide sequences that can be used as a background sequence set for comparison, and has been used to evaluate co-evolution of RNA processing signals within coding sequence [69].

To assess the representation of 5’ ss and branch point sequences, we chose two 6-mers as proxies – one for the 5’ ss (GTATGT), and one for the branch point (ACTAAC). We counted the number of times each appeared in the extant S. cerevisiae ORF set, and in each of the 10,000 permuted ORF sets, and plotted them. For both 6-mers, their counts in the extant ORF set are less than 3 standard deviations below the mean of their respective counts from the 10,000 permuted ORF sets (vertical lines), indicating that these 6-mers are significantly underrepresented in the natural *S. cerevisiae* coding sequences as compared to the randomized coding sequences (Fig 3A).

**Figure 3.**
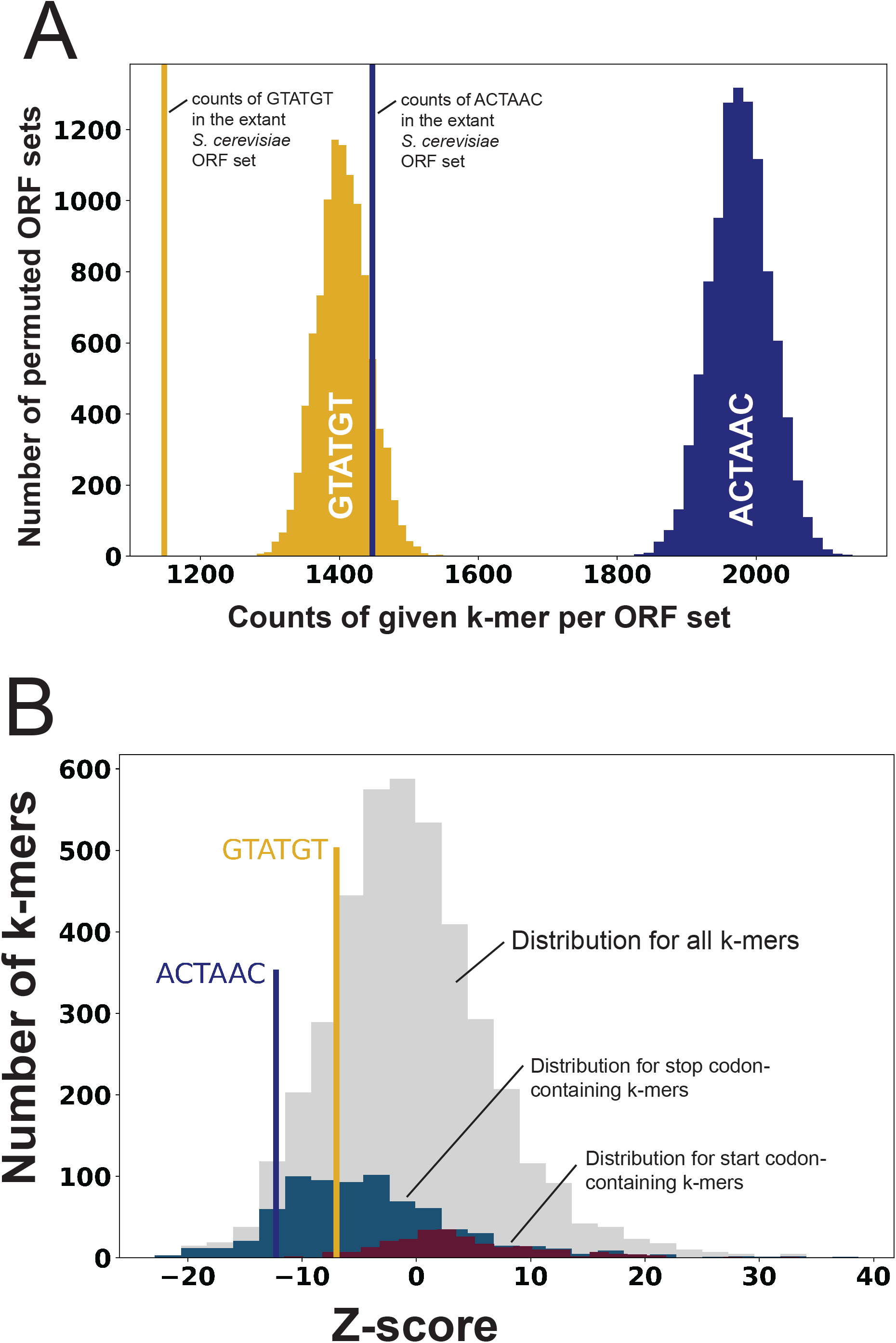
Hexamers representing splicing signals are depleted from annotated *S. cerevisiae* ORFs. (**A**) Histogram of counts of two hexamers (6-mers) serving as proxies for the branch point (ACTAAC, blue) and the 5’ ss (GTATGT, yellow) in the extant ORF set of *S. cerevisiae* (vertical lines) as compared with the distribution of counts in each of 10,000 codon-permuted ORF sets. (**B)** Histogram of Z-scores computed for each of 4096 6-mers in the extant *S. cerevisiae* ORF set relative to their corresponding mean representation in 10,000 codon-permuted *S. cerevisiae* ORF sets. The number of 6-mers (y-axis) with the given Z-score (x-axis) is represented as a histogram in grey. Similar distributions are shown for two subclasses: those containing stop codons (blue histogram) and those containing start codons (maroon histogram). The Z-scores for the branchpoint proxy hexamer ACTAAC and the 5’ splice proxy hexamer GTATGT are marked in the plot. The 6-mer “ACTAAC” had a Z-score of −12.25 and ranked 153^rd^ lowest among all 4096 6-mers, and 91^st^ lowest of 759 6-mers carrying stop codons. The 6-mer GTATGT had a Z-score of −6.98 and ranked 671^st^ lowest among all 4096 6-mers, and 7^th^ lowest of 255 6-mers carrying start codons.

To determine whether these proxy 6-mers were unusually depleted as compared to other 6-mers, we calculated a Z-score for each of the 4096 6-mer sequences in turn, comparing the counts of each in the extant *S. cerevisiae* ORF set with its mean counts in the 10,000 permuted ORF sets, and plotted the distribution (Fig 3B, grey bars). The intron branchpoint 6-mer “ACTAAC” is found comparatively much less frequently in the extant genome than are other 6-mers (Fig 3B), and much less frequently than the average stop codon-containing 6-mer (blue bars). The 5’ ss 6-mer “GTATGT” is also more depleted than the average 6-mer, especially when compared to the subset of ATG containing 6-mers (maroon bars, Fig 3B). Because there is a large amount of information in coding sequences, we cannot be certain the depletion of the splicing signal 6-mers is due to splicing. Numerous other features are being randomized by the process of codon permutation, for example 6-mer representation may be influenced by di-codon frequencies that affect translation [70]. Even so, these observations are consistent with the idea that splicing signals within ORFs carry a risk of reduced fitness. This suggests that a robust level of spliceosome activity may be sufficient to lead to loss of correct mRNA should genetic drift create splice sites within ORFs, providing a rationale for tight regulation of a splicing activity limited to pre-mRNAs that can compete [28].

### Protointrons are idiosyncratic to closely related species

*S. cerevisiae* protointrons are not conserved in closely related yeasts, suggesting that they have appeared or disappeared in *S. cerevisiae* in the time since these lineages diverged. To explore the hypothesis that protointrons arise and disappear differently in the other *Saccharomyces* lineages, we treated cultures of *S. mikatae* and *S. bayanus* with rapamycin for 0 or 60 minutes, isolated RNA, depleted rRNA, and made cDNA libraries for sequencing. These strains are NMD competent (*UPF1*), so our ability to detect transcripts subject to NMD is limited. Regardless, nearly all annotated introns in *S. cerevisiae* are present in the *S. mikatae* and *S. bayanus* genomes, and *S. bayanus* has an additional standard intron in a *CHA4*-like gene that has no ortholog in *S. cerevisiae* (Tables S6 and S7). Some annotated *S. cerevisiae* introns are missing in these close relatives and based in part on their locations and splicing efficiencies, we propose reclassifying them as protointrons (see below). In contrast to the high conservation of standard intron locations, we detect distinct sets of protointrons in each species (Tables S6 and S7). We validated a subset of these (Fig 4) and find that most all of the *S. mikatae* protointrons are not present in either *S. cerevisiae* or *S. bayanus*, and that the *S. bayanus* protointrons are not found in *S. cerevisiae* or *S. mikatae* (Tables S3, S6, and S7). An exception to this is *YIL048W*/*NEO1*, in which the same protointron is observed in both *S. cerevisiae* (Table S3) and *S. mikatae* (Fig 4, Table S6). High sequence conservation in the coding region of *YIL048W/NEO1*, most likely due to functional constraints on the protein coding function of the sequence, has fortuitously preserved the splicing signals in all of the *Saccharomyces* yeasts. The intron is in frame with the coding sequence (Table S8), thus although the splicing of this protointron is not efficient, it remains possible this intron could generate a functional alternative protein. We conclude that protointrons are idiosyncratic in closely related yeast species. This is evidence for rapid evolutionary appearance and disappearance of sequences that can be functionally recognized by the spliceosome. The dynamics of creation of protointrons thus appears consistent with genetic drift, primarily in the rapidly evolving nonconserved sequences of recently diverged genomes.

**Figure 4.**
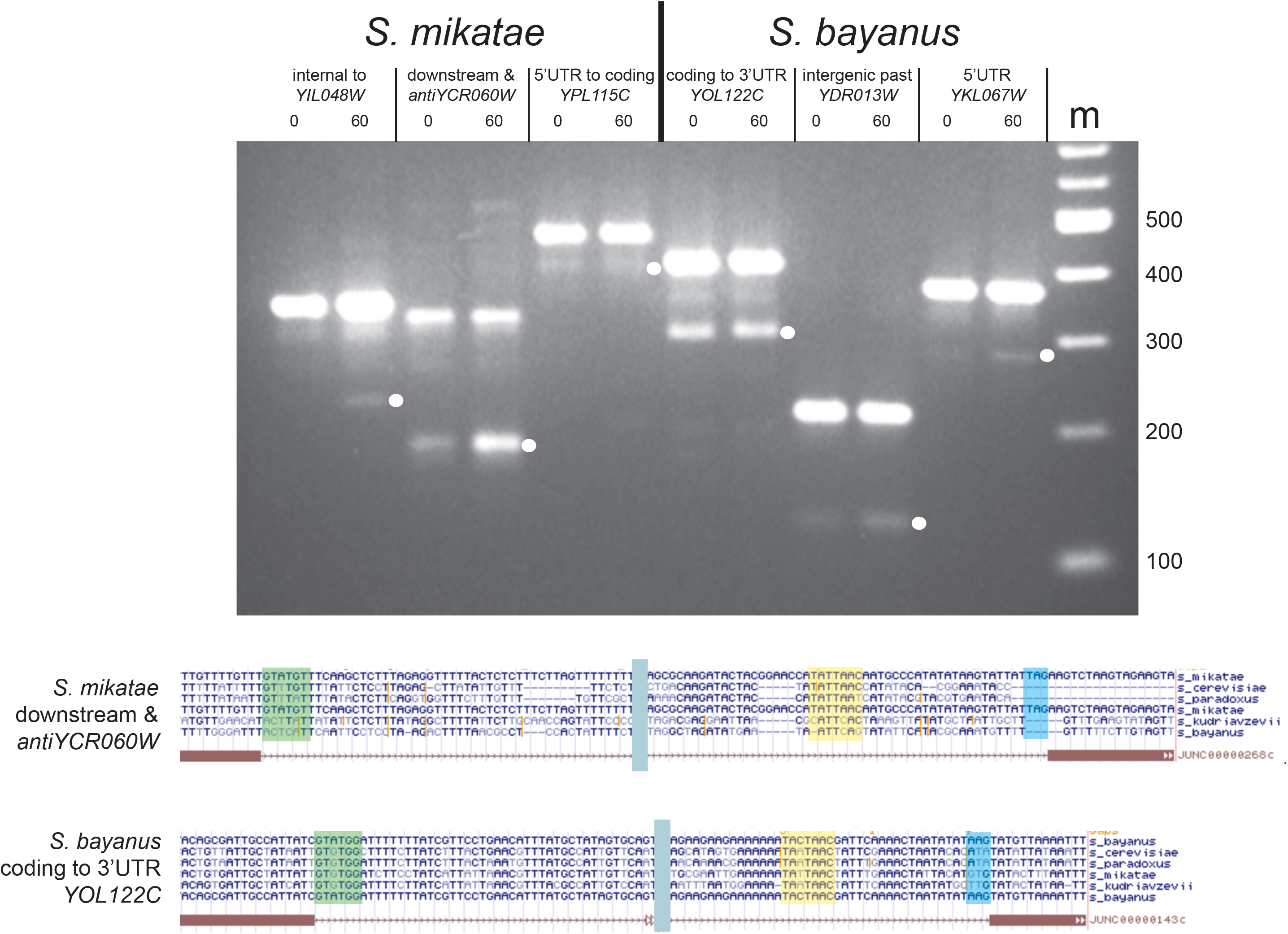
Protointrons are found in other *Saccharomyces* species but are not conserved. RT-PCR products from RNA of *S. mikatae* (left) and *S. bayanus* (right) at different protointrons identified by RNAseq. Splice junctions were validated by cloning and sequencing the PCR products indicated by a white dot. Below the gel image are shown alignments of the RT-PCR product sequences from *S. mikatae antiYCR060W* and *S. bayanus YOL122C* to their corresponding genomes to show lack of conservation of splicing signals (boxed).

### Introns with features of both protointrons and standard introns may be intermediates in de novo intron creation

Based on the above analysis and those described elsewhere that note “novel” splicing [32-34, 37-39, 67], we propose defining standard introns as (1) conserved in related organisms or clades, (2) efficiently spliced under appropriate physiological conditions, and (3) established in the pathway for production or regulation of a functional gene product. Furthermore, we propose defining protointrons as (1) not conserved in closely related species, (2) inefficiently spliced, and (3) not clearly understood to contribute to correct expression or regulation of a gene. This simple definition allows classification of most all of the observed splicing events in the yeast genome as either protointron or standard intron, with only a few exceptions. The vast majority of protointrons arise by neutral drift and likely provide no fitness advantage, and most probably disappear. A few introns do not neatly fall into one or the other category and may be transitioning from protointron to standard intron status, as predicted by the intronization model. Like protointrons these intermediate class introns not conserved, but unlike typical protointrons they have increased splicing efficiency, and may appear positioned to influence expression of the gene that carries them.

Examples of protointrons that may be on an evolutionary path toward standard intron status include introns in the 5’ UTRs of *S. cerevisiae MTR2, USV1, YEL023C*, and *MCR1*, and in the 5’ UTR of *S. bayanus YTA12* (Fig 5). Using the unrooted tree describing relationships between the genomes of the sensu stricto yeasts [71], we map the appearance of these high efficiency protointrons as predicted by their presence in extant genomes that have diverged over 10-20 million years. The *MTR2* intron contains essential sequences [72] and can diversify the N-terminal sequence of the mRNA export protein Mtr2 [73]. When sequences of related yeasts became available [31], it became clear that the *MTR2* intron is unique to *S. cerevisiae* (Fig 5). High efficiency protointrons are found in the 5’UTRs of *USV1* and *MCR1*. The *USV1* intron is efficiently spliced after rapamycin treatment in *S. cerevisiae* and is also functional in *S. mikatae* (Fig 5, Table S6). However, *S. kudiravzevii* and *S. bayanus* have different sets of nucleotide changes that eliminate splice sites and branchpoints required for this intron. The *MCR1* intron is spliced at about 30%, and appears to be shared by *S. paradoxus*, but is absent in *S. mikatae*, *S. bayanus*, and *S. kudriavzevii*. None of these splicing events alter the N-terminus of Usv1 (a stress induced transcription factor) or Mcr1 (a mitochondrial NADH-cytochrome b5 reductase), however both introns remove uORFs from the 5’ UTRs of these genes, suggesting that splicing could affect 5’UTR function in mRNA translation or stability for both genes. Finally, a very efficiently spliced (>95%) intron is found in the 5’UTR of the *S. bayanus YTA12* gene, as well as in *S. kudriavzevii* (Fig 5). Removal of this intron does not alter the N-terminus of Yta12, a mitochondrial protein complex assembly factor [74], but does lead to removal of a uORF. Interestingly the *S. cerevisiae YTA12* gene matches at 88 of 117 (75%) positions in the *S. bayanus* intron and has neither an intron nor any uORF (see below).

**Figure 5.**
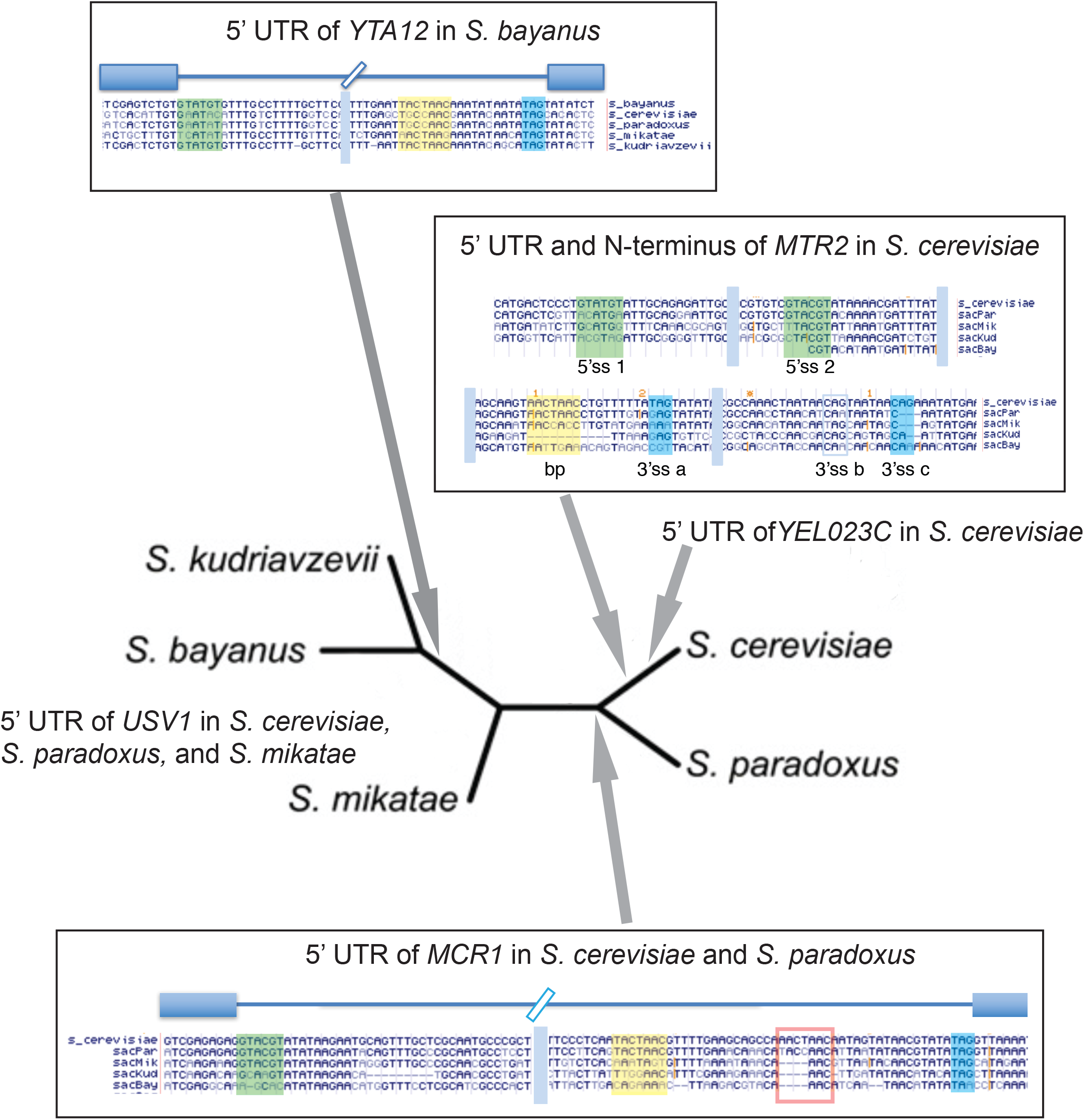
Unusually efficient protointrons that may be evolving toward standard introns. Positions of 5 efficiently spliced protointrons that share similarity with standard introns on the unrooted tree of sensu stricto *Saccharomyces* species are shown. Grey arrows indicate separation points that delineate boundaries between species having or lacking the indicated protointron sequence. Bars in the alignments indicate that sequences between these blocks are not shown. 5’ ss are green, branchpoint sequences are yellow, and 3’ ss are blue. Although these protointrons are restricted to one or two closely related species, their splicing efficiency approaches that of standard introns. Most protointrons are unique to a species and are very inefficiently spliced.

Ten percent of the protointrons identified in *S. cerevisiae* are located exclusively in 5’UTRs (Supplemental Figure S1B). The apparent relationship between introns and uORFs leads to the idea that 5’UTR introns may be adaptive by protecting mRNAs with long 5’UTRs from the general negative effect of uORFs [48, 75]. To test this idea, we asked whether uORFs are present more frequently in 5’UTRs that contain standard introns, as compared to similarly sized 5’UTRs that do not. There are 22 yeast genes (7%) with standard introns in their 5’UTRs, (this number does not count the annotated introns in *MCR1*, *MTR2*, or *USV1*, which are not conserved across the *sensu stricto* group). The size range of the (unspliced) 5’UTRs for these 22 is from ~240 to 950 nucleotides, and there are 91 intronless genes with 5’UTRs in this size range. We counted uORFs longer than 4 codons (including the AUG, but not the stop codon) within 5’UTRs in this size range. Among the 91 genes without 5’ UTR introns, 53 lack any uORFs, whereas 38 have at least one uORF. All 22 genes with 5’UTR introns have at least one uORF, and for 20 of these all the uORFs in the 5’ UTR are removed by splicing. This distribution of uORFs in 5’UTRs with introns is unlikely to have been generated by chance (Fisher’s exact test, p <10^-5^). One hypothesis to explain this is that by removing much of the 5’ UTR RNA, an intron may protect a gene that has a long 5’UTR from genetic drift that creates uORFs, or other translational inhibitory features like RNA secondary structure [48]. Additional experiments will be needed to determine which if any of these splicing events promotes gene expression and whether or not the effect contributes to fitness.

A second evolutionary scenario whereby protointrons may be adaptive concerns in frame splicing, which would produce an alternative polypeptide. This appears to be the case for the standard intron in *PTC7* [30, 66]. A similar intron is found in *MRM2*, where as many as three different proteins may be produced (Tables S1 and S2, see also [37], NB: not annotated in SGD, but this fits the standard intron definition). Despite forces that seem to deplete splicing signals within ORFs (Fig 3), we find 20 (out of 63) 3n protointrons within ORFs that do not interrupt the reading frame and thus could produce functional proteins, particularly under stress or other conditions where RPG transcription is reduced (Table S8). We suggest that such protointrons provide evolutionary opportunities to create new protein isoforms from existing genes.

### Telomeric Y’ repeats

Intron-like sequences have been noted in the telomeric Y’ family of repeat sequences for more than 25 years and continue to be annotated in the Saccharomyces Genome Database (SGD). So far, molecular tests for splicing of these annotated introns have been negative [61, 76]. Intron predictions allow some Y’ repeat element copies to encode a large (1838 amino acid) protein (e. g. YNL339C, Fig 6A), that carries an N-terminal Sir1 domain and a central DExD helicase domain. Other Y’ elements differ in sequence and can only encode fragments of the open reading frame that may nonetheless produce smaller functional proteins, for example the helicase overexpressed in telomerase-deficient “escaper” colonies [76, 77]. The function(s) of any of the Y’ element predicted proteins in normal cells are not known.

**Figure 6.**
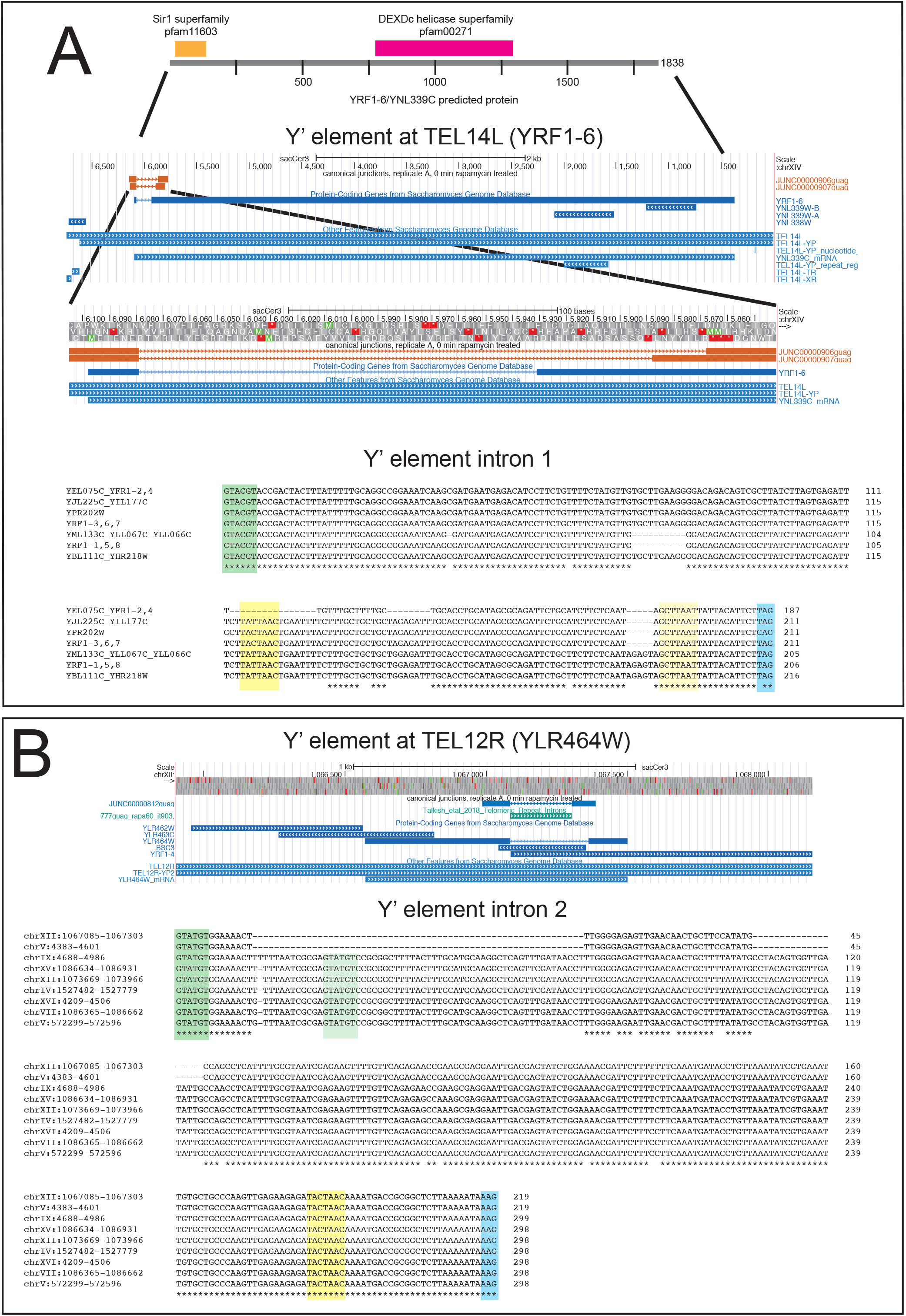
Introns in the Y’ element repeat family. Two different introns are found in the transcribed Y’ repeat elements. **(A)** Y’ intron 1. Top: expanded view of the protein encoded by *YRF1-6* located in the Y’ element at the left end of chromosome XIV with the Sir1 and DECD helicase homology regions indicated. An expanded segment from the upstream part of the gene shows the alignment of the detected intron relative to the annotated predicted intron at SGD. At the bottom is shown the alignment of the seven different versions of this intron from the seventeen Y’ elements in the *S. cerevisiae* genome that possess it. Sequence names are based on standard and systematic annotations from the *Saccharomyces* Genome Database (SGD). 5’ ss are green, branchpoint sequences are yellow, and 3’ ss are blue. **(B)** Y’ intron 2. Top: expanded view of the protein encoded by *YLR464W* located in the Y’ element at the right end of chromosome XII. The alignment shows the detected intron relative to the annotated predicted intron at SGD. At the bottom is shown the alignment of the nine different versions of this intron from the nine Y’ elements in the *S. cerevisiae* genome that possess it. Splicing signals are highlighted as in (A).

To determine if Y’ element transcripts are spliced, we allowed RNAseq reads to map to the repetitive Y’ elements (i. e. without masking). Although the mapped locations may not be the precise origin of the RNA that created the read, this allows us to identify spliced reads and assign them to possible members of the Y’ repeat family. We find two introns within the Y’ repeat family (Fig 6), one of which lies on the far left of the repeat and is required to create the open reading frame for the longest predicted protein (exemplified by YNL339C near TEL14L, Fig 6A). The other is in the center of the Y’ repeat (exemplified by YLR464W near TEL12R, Fig 6B). Neither of these introns matches the annotations at SGD, and instead in both cases, downstream 3’ ss are used (see also [37]). It is not uncommon for the yeast spliceosome to skip proximal 3’ ss in favor of a distal 3’ ss, in some cases due to secondary structure of the pre-mRNA [78]. The Y’ elements in *S. cerevisiae* differ from each other; not all can express a protein as large as YNL399C after removal of intron 1 using the distal 3’ ss. Intron 2 splicing does not greatly extend the open reading frame of YLR464W. To confirm that the reads arise from splicing rather than from a deleted copy of the Y’ element precisely lacking the intron, we searched the genome using the “spliced” sequence produced for intron 1 or intron 2 and found that there is no such contiguous genomic sequence. We conclude that Y’ element transcripts can carry at least two introns that are distinct from current annotations in SGD (Fig 6).

To evaluate the sequence relationships of the Y’ repeat element introns we aligned them with each other, after merging identical copies into one. All the predicted intron 1 sequences have the second most common 5’ ss in the yeast genome (GUACGU, followed by the preferred A at position 7, [61]. The most distal 3’ ss of several possible creates the large ORF, and is UAG for all except YPR202W, which has a CAG. Several other potential 3’ ss (including the one annotated in SGD) are skipped or used alternatively. Interestingly only YPR202W, YRF1-3, YRF1-6, and YRF1-7 have the canonical UACUAAC branch point sequence, whereas most of the others have UAUUAAC, a variant found in some standard introns (Fig 6A). The remaining group (YEL075C, YRF1-2, and YRF1-4) are deleted for the region containing the branch point, suggesting that they are unable to be spliced. Intron 2 has the most common 5’ss GUAUGU, and the most common branch point UACUAAC, and uses the first AAG (a less commonly used but standard 3’ ss) downstream from the branch site (Fig 6B). Most copies also contain an alternative 5’ ss which is used less frequently. We have not estimated the efficiency of splicing of these introns because we cannot reliably assign reads to specific repeat elements with confidence. The current genome assemblies of *S. mikatae*, *S. paradoxus*, and *S. kudriavzevii*, but not the *S. bayanus* assembly include at least one Y’ element related to the *S. cerevisiae* elements [71], but the precise numbers and arrangements of the Y’ elements in those genomes await refinement of the genome assemblies for those organisms.

### The *S. bayanus* YTA12 protointron functions in *S. cerevisiae*

The finding of a highly efficient protointron in the *YTA12* 5’ UTR of *S. bayanus* (and putatively in *S. kudriavzevii,* Fig 5) that is not observed in the alignable syntenic sequence of *S. cerevisiae* prompted us to test (1) whether this *S. bayanus*-specific intron can be spliced in *S. cerevisiae*, and (2) whether the intron might confer some advantage for growth on glycerol, given the function of Yta12 in assembly of mitochondrial protein complexes [74]. Fig 7A shows an alignment of the region including and upstream of the Yta12 start codon from *S. bayanus* (sacBay), *S. cerevisiae* (sacCer) and *S. cerevisiae* in which the 117 bp of the *S. cerevisiae* genome corresponding to the *S. bayanus* intron have been replaced in *S. cerevisiae* with the *S. bayanus* intron (Sc-SbI). This replacement was made using CRISPR/Cas9 guided cleavage of an *S. cerevisiae*-specific target sequence within the syntenic region and a repair fragment containing the *S. bayanus* intron (Fig 7B). As controls, we created *S. cerevisiae* strains precisely deleted for the syntenic region aligning with the *S. bayanus* intron, as well as versions of the *S. bayanus* intron with 5’ ss mutations (Fig 7C). We isolated RNA and evaluated the expression of these modified *YTA12* genes by extension of a labeled primer complementary to *YTA12* mRNA with reverse transcriptase (Fig 7C). The major transcription start sites for *YTA12* in *S. cerevisiae* map about 300 nt from the 5’ end of the primer (Fig 7A and C, lane 1). These start sites are unaffected by the 117 bp deletion (the same collection of cDNAs are shorter by 117 residues, lane 2). The migration of cDNAs from the deletion strain are useful to mark the expected position of spliced RNAs, and indeed replacement of the 117 bp with the *S. bayanus* intron sequence (lane 3) results in the appearance of the same collection of cDNAs with the disappearance of the signal from pre-mRNA, indicating efficient splicing (lane 3, compare with lane 1).

**Figure 7.**
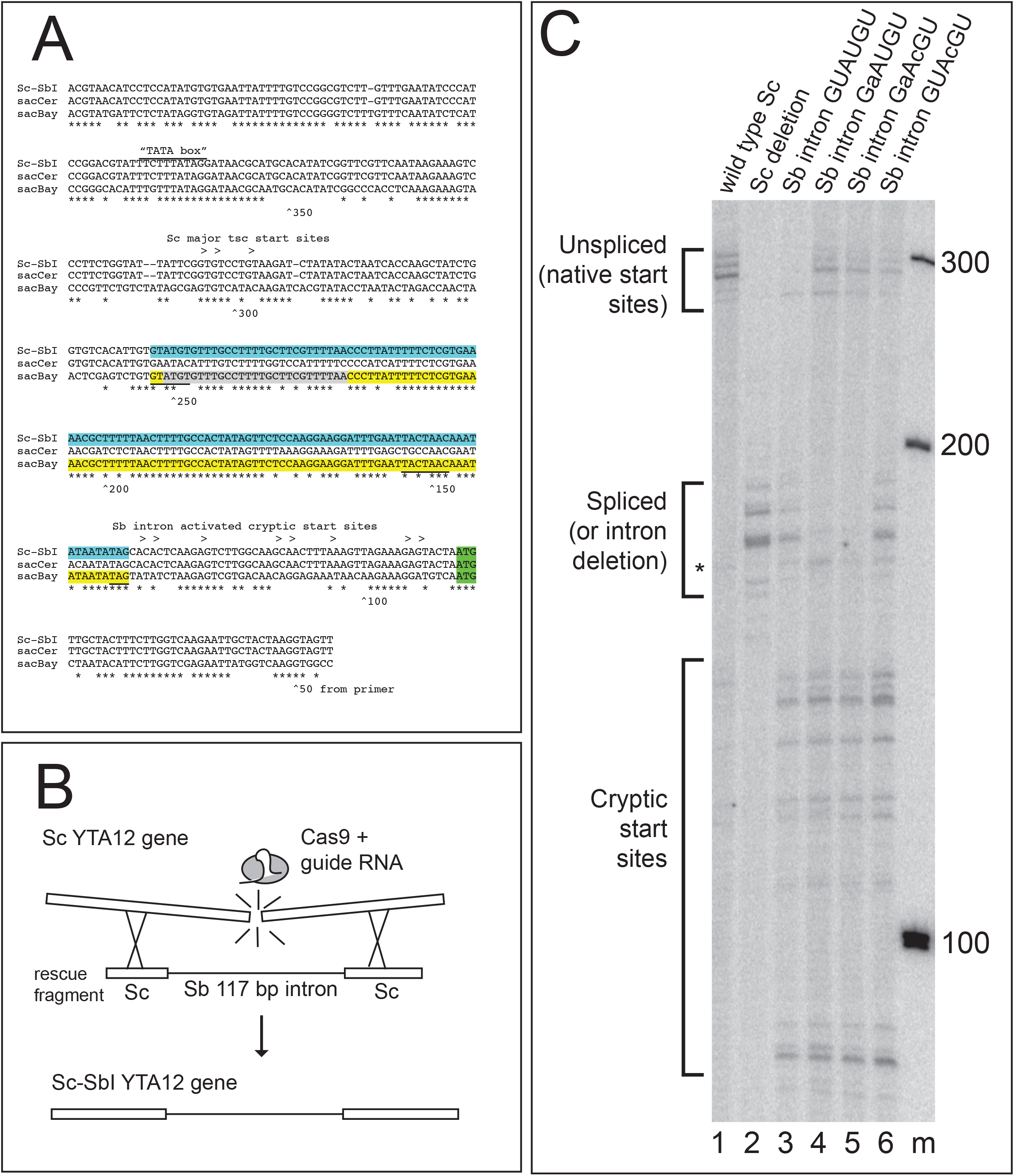
The *S. bayanus* YTA12 5’UTR intron is efficiently spliced in *S. cerevisiae*. **(A)** Alignment of the *YTA12* promoter and 5’UTR from *S.cerevisiae* (sacCer, no intron), *S. bayanus* (sacBay, very efficient intron), and the *S. cerevisiae* strain carrying the *S. bayanus* intron (Sc-SbI), showing the major transcription start sites and the cryptic start sites (>), the splice sites (underlined), and the aligned base pairs (*). **(B)** Strategy for CRISPR/Cas9 editing-based transplantation of the *S. bayanus* intron into *S. cerevisiae*. A guide sequence was designed to recognize a sequence present in the *S. cerevisiae YTA12* 5’UTR but not present in the *S. bayanus* intron. A plasmid derived from those provided by DiCarlo et al. [97] expressing this guide along with Cas9 was co-transformed with a synthetic rescue fragment that contained the *S. bayanus* intron sequence between “exons” from *S. cerevisiae*. Repair of the double-stranded break using this rescue fragment results in transplantation of the *S. bayanus* intron into *S. cerevisiae*. **(C)** Reverse transcriptase primer extension analysis of RNA from the YTA12 locus of S. cerevisiae strains with the transplanted *S. bayanus* intron and mutant derivatives. The cDNAs representing unspliced (native) start sites, spliced (or deleted) RNAs initiating from the normal start site, and unspliced RNAs arising from cryptic start sites are indicated at left. Lane 1, wild type; lane 2, deletion of the region that aligns with the *S. bayanus* intron; lane 3, transplantation of the wild type S. bayanus intron; lane 4, GaAUGU mutation of the *S. bayanus* intron 5’ ss; lane 5, GaAcGU mutant eliminating both the 5’ ss and the start codon of a uORF; lane 6, GUAcGU mutant that creates a common functional ss while removing the start codon of the uORF; lane m, 100 bp ladder markers.

Mutation of the 5’ ss from GUAUGU to GaAUGU or GaAcGU results in the reduction of the spliced mRNA cDNAs, and the appearance of cDNAs corresponding to pre-mRNA (lanes 4 and 5), indicating that splicing is inhibited by these mutations. Changing the 5’ ss from GUAUGU to the less commonly used GUAcGU reduces the efficiency of splicing but does not block it, as judged by the slight accumulation of unspliced RNA (lane 6). Unexpectedly, the *S. bayanus* intron sequence activates a set of cryptic start sites in the *S. cerevisiae* sequences downstream of the major start site and the *S. bayanus* intron (Fig 7C, lanes 2-6). These start sites are inefficiently used in the wild type *S. cerevisiae YTA12* promoter (lane 1). One consequence is that new mRNAs are made that initiate downstream of the intron and thus do not require splicing for expression of Yta12. This interpretation is supported by the observation that all the strains grow on YP glycerol plates as well as wild type BY4741 (not shown). This result highlights the challenge of anticipating the effect of mutations in 5’ UTRs where transcription, splicing, and translation operate together on the same sequence. This experiment measures changes in splicing due only to differences in the intron, and not due to any differences in exonic sequences or trans-acting factors between *S. bayanus* and *S. cerevisiae*. We conclude that the efficiently spliced protointron from *S. bayanus* is equally at home in *S. cerevisiae*. This intron appears to have formed in *S. bayanus* after *S. bayanus* and *S. cerevisiae* last shared a common ancestor, but before the divergence of *S. bayanus* from *S. kudriavzevii*.

## Discussion

### A second class of splicing events exposes roles of the spliceosome in evolution

Here we characterize a class of splicing events in yeast we call protointrons. Many previous studies have noted “novel” introns in yeast under a variety of experimental conditions and genetic backgrounds [32-39, 67]. Here we distinguish protointrons by several criteria, most importantly that they reside at locations not overlapping known standard introns. We first recognized protointrons while studying how the abrupt disappearance of RPG pre-mRNA during early nutrient deprivation signaling frees the spliceosome to increase splicing of other pre-mRNAs [28, 29]. RNAseq analysis of NMD-deficient yeast cells treated with rapamycin revealed that protointrons are found throughout the transcriptome in both coding and non-coding regions of pre-mRNAs, ncRNAs, and antisense transcripts, such as CUTs, SUTs, and XUTs (Fig 1, Fig S1). Protointrons contain all of the splicing signals necessary for recognition by the spliceosome (5’SS, BP, and 3’SS), however the sequences of these signals are more variable than those of standard introns (Fig 2A and B). Whereas standard introns are conserved in related organisms, efficiently spliced, and established for production or regulation of a functional gene product, protointrons are present in one or a few closely related species, not efficiently spliced, and do not clearly contribute to correct expression or regulation of a gene. Given this redefinition, we propose a revised intron annotation, including the addition of a standard intron in *MRM2*, and molecular evidence for the correct location of expected but not demonstrated splicing of Y’ repeat element transcripts. We provide this and related data on a publicly accessible genome browser with several *Saccharomyces* species genomes at http://intron.ucsc.edu/.

Splicing events that occur outside our expectation of what is needed to make a protein or a structural RNA have attracted labels like “splicing noise” or “splicing error” [79]. But viewing the spliceosome as an enzyme able to catalyze a complex series of pre-mRNA binding, refolding, and release operations, including two cleavage-ligation reactions, or even just the first one [35, 80], on a very diverse set of substrates (for review see [2]) suggests that such terms should be more carefully defined. The protointrons described here, as well as for example similar newly evolved splicing events observed in mammalian lncRNAs [81, 82] reveal the outer edges of the substrate repertoire of this enzyme in sequence space, and do not represent either splicing noise or splicing errors. We suggest the term “splicing noise” should refer to fluctuations due to stochastic events affecting particular splicing events, just as the term “transcriptional noise” refers to the stochasticity of transcription events (see [83] and references therein). We also suggest the term “splicing error” should refer to events within the spliceosome that lead to spliceosome assembly or catalysis that is incompatible with successful completion of the two splicing reactions, spliced product release, and recycling. In order for splicing to contribute to rapid evolution of multicellular organisms it seems likely that a variety of sequences besides highly evolved introns would need to be recognized and spliced, including those that appear in genomes by genetic drift. The extent to which these spliceosome-generated spliced RNA sequences contribute to fitness would eventually determine their evolutionary fate. The protointron class of splicing substrates represents opportunity to create new genes, create new proteins from existing genes, or impose new regulatory controls on existing genes.

### Some protointrons show greater splicing efficiency and may be adaptive

The forces and mechanisms that drive intron evolution in eukaryotic genomes are still largely unknown. If protointrons represent raw material for intron creation by the process of intronization, then perhaps the most efficiently spliced protointrons represent intermediates in standard intron formation that are advancing by selection of improving mutations. Our data provide evidence for rapid and complete intronization in the *YTA12* 5’ UTR between now and the time *S. bayanus* and *S. cerevisiae* last shared a common ancestor (~ 20 Mya, [71]). Over the 117 bp intron sequence, the *S. cerevisiae* 5’ UTR differs at 29 positions (Fig 7A). Replacement of this region with the *S. bayanus* sequence produces an efficiently spliced intron in *S. cerevisiae* (Fig 7C). This intron transplantation experiment shows that no species-specific barrier prevents this sequence from serving as an efficient intron in *S. cerevisiae*. Although this intron appears fixed in the *S. bayanus* and *S. kudravzevii* branch of the *Saccharomyces* tree, there is currently no evidence for fitness effects, and thus this intron could be a product of neutral evolution.

In some cases, a protointron might provide increased fitness that would explain its evolutionary persistence. We suggest three specific ways that protointrons may support improvements in gene function. Approximately 10% of protointrons reside entirely within 5’UTRs (Supplemental Figure S1B), including the four most efficiently spliced protointrons we observed (*S. cerevisiae MTR2*, *USV1*, and *MCR1*, *S. bayanus YTA12*). We realized that genes with long distances between their transcription start sites and their start codons (i. e. with large 5’ UTRs) are at risk for mutations that create a uORF in the 5’ UTR, which often negatively influences translation [48, 75]. Removal of a large region of the UTR by splicing would buffer this genetic risk. Secondary structures or other detrimental sequences that might arise in long 5’UTRs [84] might also be safely removed by splicing. To test the plausibility of this idea, we examined the frequency of uORFs in 5’ UTRs of *S. cerevisiae* genes that have or do not have 5’ UTR introns and found that uORFs are significantly more prevalent in 5’ UTRs that have introns as compared to other 5’ UTRs (see Results). This suggests that the presence of a 5’ UTR intron may help buffer an mRNA against any detrimental effects of uORFs or RNA secondary structure, and provides evidence that intronization in particular in 5’ UTRs may be adaptive in *Saccharomyces* species.

A second way that protointrons may become functional is by producing in frame splicing events within ORFs to create mRNAs encoding shorter protein isoforms with new functions. The frequency of splicing signals is lower than expected in *S. cerevisiae* ORFs (Fig 3), supporting the expectation that most introns that arise within ORFs would be detrimental to fitness. Despite this, we found 20 protointrons contained within ORFs that do not interrupt the reading frame, and that may lead to the translation of alternative protein products (Table S8). If such shorter proteins contribute to fitness, mutations that increase the splicing of the protointron (without disrupting the function of the full-length protein) may lead to the establishment and conservation of a standard intron that allows production of both protein forms. This may be the mechanism by which the conserved in-frame introns of *PTC7* [30, 66] and *MRM2* ([37], this work) have evolved. In these cases, both the intron and the protein sequence encoded by the intron are conserved in *sensu stricto* yeasts, suggesting both contribute to fitness across the genus *Saccharomyces*. Many protointrons span the boundaries between conserved and nonconserved sequences (Fig 2B), increasing the chances that a new splicing event will alter one or the other end of an existing protein. Studies of protein evolution indicate that proteins evolve at their edges [85], suggesting that protointrons may contribute to this as well. Although there is as yet no evidence for new function, the 5’ UTR protointron in the *S. cerevisiae MTR2* gene has arisen sufficiently close to the start codon that different alternative splicing events add different peptides to the amino terminus of the annotated protein sequence [72, 73]. Thus, protointrons that appear in frame within existing genes, or that span the edges of existing genes, create protein expression variation that may provide fitness advantages, in particular under stress conditions that have yet to be explored.

A third way that protointrons may prove advantageous is through controlled downregulation through splicing and NMD. We find that 16% of protointrons in *S. cerevisiae* span the 5’ UTR and coding region of twenty-seven genes and upon being spliced, remove the canonical AUG start codon making these transcripts potential targets of NMD. In ten of these protointrons, the AUG start codon is embedded within the GU**AUG** of the 5’ ss (e. g. Ade2), suggesting sequences surrounding start codons are particularly susceptible to drifting toward a 5’ ss. A recently studied example of this is the standard intron in the *PRP5* gene that is conserved in the *Saccharomyces* genus and destroys the *PRP5* mRNA by removing the start codon and creating a transcript that is subject to NMD [65]. The intron must have appeared since the divergence of the *Saccharomyces* species from their common ancestor with *Lachancia kluveri*, since this more distant relative has a different intron in its *PRP5* gene. This situation may evolve where overexpression of a particular protein may be detrimental. *PRP5* encodes a splicing factor, and increase (or decrease) in Prp5 protein activity may increase splicing and reduce (or decrease splicing and accumulate) *PRP5* mRNA levels by using this conserved out of frame intron to create a homeostatic regulatory loop. A more difficult to recognize but no less important mechanism is illustrated by the *BDF2* gene in which abortive splicing downregulates expression through spliceosome-mediated decay [35]. It is unclear whether to annotate this location and others like it [80] as an intron, since it does not appear that 3’ splice site selection is required or important for its activity. Thus, even protointrons that are out of frame within coding regions, or pseudo-intron locations at which abortive splicing takes place may provide opportunities for adaptive regulatory controls to evolve.

### Conclusions

Protointrons are a rare class of splicing events that represent the action of the spliceosome on RNA without a necessary connection to the expression of a mature gene. In mammalian cells the spliceosome is no less constrained, and a very large number of alternative splicing events that appear unrelated to “correct” gene expression support this [86]. In particular, newly evolved lncRNAs have introns that are inefficiently spliced and have multiple alternative splice sites, unlike older, more conserved lncRNA and mRNA encoding genes [81, 82]. These observations indicate that a general feature of the evolution of introns is that any transcribed sequence has a chance of being spliced by the spliceosome, should that sequence evolve recognizable splicing signals. Additionally, any sequence that suddenly becomes transcribed can be expected to contain sequences by chance that are immediately recognized as introns. Since the sequences required for splicing are ubiquitous and have low information, many such newly appearing sequences will immediately produce diverse RNA transcripts. If these confer some advantage, or if mutations that improve splicing become fixed by neutral genetic drift, then a standard intron may evolve. This general pathway may be a source of new introns whose splicing contributes to diversification of the transcriptome, and to the appearance of new genes and new products from existing genes, as genomes evolve.

## Materials and Methods

### Strains and culture conditions

Two independent cultures of *S. cerevisiae* strain BY4741 *upf1Δ* (*MATa his3Δ1 leu2Δ0 met15Δ0 ura3Δ0 upf1Δ::KANMX*) were grown in YEPD medium at 30°C to an optical density at 600 nm (OD_600_) ≈ 0.5. The cultures were split and rapamycin was added to one half at a final concentration of 200 ng/ml for 1 hour. *S. bayanus* strain JRY9195 (*MATa hoD::loxP his3 lys2 ura3*) and *S. mikatae* strain JRY9184 (*MATa hoD::NatMX trp1D::HygMX ura3D::HygMX*) were grown in YEPD medium at 26 °C, and were treated with rapamycin as for *S. cerevisiae* except at 26°C. These strains were a kind gift of Chris Hettinger [71].

### RNA isolation

RNA was extracted from yeast cells using Procedure 1 as described [87]. Prior to RNAseq library construction (see below), RNA was DNased using Turbo DNase (Life Technologies) and RNA quality was evaluated using the 2100 Bioanalyzer (Agilent).

### RNAseq library preparation

5-10 ug of total *S. cerevisiae* RNA was depleted of ribosomal RNA using the RiboZero Gold rRNA Removal Kit (Illumina) according to the manufacturer’s instructions. Strand-specific cDNA libraries were prepared using the Kapa Stranded RNA-Seq Library Preparation Kit for Illumina Platforms (Kapa Biosciences) following the manufacturer’s instructions with the following modifications. Sequencing adapters and oligonucleotides used for PCR barcoding were from the NEBNext Multiplex Oligos for Illumina Kit (New England Biolabs, NEB). Prior to PCR amplification of the library, adapter-ligated cDNA was treated with USER enzyme (NEB). Adapter-ligated libraries were then PCR amplified for 10 cycles using NEB index primers compatible with Illumina sequencing. After amplification, size selection of the libraries was performed using an E-gel Safe Imager and 2% E-gel size select gels (Invitrogen). Indexed libraries were pooled and 100 bp paired-end sequenced on the same flow cell of an Illumina HiSeq4000 instrument at the Berkeley sequencing facility. RNA extracted from *S. bayanus and S. mikatae* was depleted of ribosomal RNA as described above. Strand-specific cDNA sequencing libraries were prepared as described [88] and 50 bp paired-end sequenced on the HiSeq2000 platform (Illumina).

### Mapping and Analysis of RNAseq Data

RNAseq data is deposited in GEO under the accession number GSE102615. For *S. cerevisiae* libraries, all mappings were done using 100×100 bp reads to the SacCer3 Apr. 2011 genome assembly (Saccharomyces Genome Database, SGD, [89]). For *S. bayanus* and *S. mikatae* libraries, mappings were done using 51×51 bp reads to the SacBay2 and SacMik2 genome assemblies (Saccharomyces Sensu Stricto Database, [71]), respectively. Reads mapping by BowTie2 [90] to *S. cerevisiae* tRNA and rRNA defined by Ensemble or to Ty elements defined by SGD were discarded, however mappings to Y’ elements were recovered. For each library, reads were remapped to their respective genomes using STAR with two-pass mode [91]. PCR duplicate reads (reads with identical positions at both ends) were discarded and reduced down to one read. Changes in gene expression upon treatment of cells with rapamycin were determined using DESeq2 [92], comparing untreated and treated cells. Splice junctions were identified by STAR mapping [91].

Splicing Indexes (ratios of splicing measurements) were calculated by comparing reads that cross the intron, reads that cross the splice junction, and reads in exon 2 in different ways. Splice junction coverage is taken as the number of reads that cross the splice junction. Intron coverage was taken as the average per nucleotide coverage across the whole intron. When introns overlapped, a minimal length intron was used such that start of the intron was the most downstream start of the overlapping introns and the end was the most upstream start of the overlapping introns. Exon2 coverage was the average coverage for 100 bases of the following exon, using the most downstream 3’ ss to define the exon. Log2-transformed ratios (Indexes) were calculated for the three comparisons: intron/exon2, splice junction/exon2, splice junction/intron. Figure 1A shows how the splice junction/intron index changes with rapamycin treatment by plotting the value of log2[splice junction-60/intron-60] – log2[splice junction-0/intron-0] for each intron in each replicate. The splicing events plotted here are for locations whose overall transcript level changes less than 2 fold, and whose junctions are supported by at least 35 reads for standard introns or at least 50 reads for protointrons. The general shift of the points to the upper right quadrant indicates increased splicing efficiency (increased junction relative to intron reads) after rapamycin treatment.

Intron splice sites and candidate branch sits were extracted for analysis using the mapped splice junctions and by choosing a best branch point using the following heuristics. The likely branch points (underlined) were identified by searching introns for the following sequences in order until a branchpoint was identified: 1. ACTA<u>A</u>, 2. RYTR<u>A</u>YR, 3. YTR<u>A</u>Y (where R = A or G, Y = C or T) constrained to be 45 or more bases away from the 5’ ss and no closer than 7 nucleotides upstream from the 3’ ss. Candidate introns not matching YTRAY were considered to have no good match to a branchpoint consensus. If multiple equally good branchpoints are identified the one closer to the 3’ ss was recorded. Details and scripts are at: <https://github.com/donoyoyo/intron_bp_generator>. To evaluate conservation, phastCons conservation scores were extracted from a window surrounding the splice site or branchpoint using data from the UCSC Human Genome Browser for *S. cerevisiae* at <http://genome.ucsc.edu>. Weblogos [93] were created using the site at https://weblogo.berkeley.edu.

### Reverse Transcription and PCR

RNA was reverse transcribed using SuperScript III (Life Technologies) according to the manufacturer’s instructions using a mixture of anchored oligo-dT (T24VN) and random hexamers as primer. Primers to validate and sequence products of splicing from protointrons by RT-PCR were designed using Primer3 [94]. PCR was performed using oligonucleotides listed in Table S5. PCR products were resolved by electrophoresis on agarose gels and visualized with ethidium bromide staining.

### Cloning and Sanger sequencing of PCR products

PCR products generated by *T. aquaticus* DNA polymerase (Taq) were cut from low melting point agarose gels and purified using Machernary-Nagel gel extraction kits, then cloned using TOPO-cloning (Invitrogen). Inserts were sequenced by Sanger sequencing at the U. C. Berkeley sequencing center. Splice junctions were identified using BLAT [95] running behind a home copy of the UCSC Genome Browser [96] publicly available at http://intron.ucsc.edu/.

### Estimation of background frequency of splicing signals in codon-permuted yeast genes

To test the hypothesis that “ACTAAC” (proxy for the branchpoint sequence), GTATGT (proxy for the 5’ ss), or any other 6-mer nucleotide sequence within extant yeast ORFs might be enriched or depleted, we created 10,000 codon-permuted versions of the *S. cerevisiae* ORF set and counted the number of each of the 4096 possible 6-mers in each, computing a Z-score for each that compares representation of each in the extant ORF set to the mean representation of each in the 10,000 permuted ORF sets. To create permuted ORF sets in a way that preserves the GC content and codon usage of the extant set, we permuted the codons within each ORF (except for the start and stop codons) in the complete set of ORFs. Scripts for creating permuted ORF sets and analysis related to this question can be found under this github link: https://github.com/rshelans/genePermuter

### CRISPR/Cas9 mediated intron transplantation

Yeast CRISPR editing was done essentially as described by DiCarlo et al [97], except that we rearranged the elements from different plasmids into a simplified single plasmid system by Gibson assembly. We obtained p426-crRNA-CAN1.Y and p414-TEF1p-Cas9-CYC1t [97] from Addgene. To create a BaeI cleavable cassette for easy guide cloning, we annealed oligos newguide1 and newguide2 together, and separately newguide3 and newguide4, and filled to make two fragments which were mixed and then PCR amplified using newguide1 and newguide4 as primers (Table S5). This duplex was purified and assembled using Gibson mix (NEB) with p426-crRNA-CAN1.Y that had been cut with NheI and Acc65I to replace the CAN1.Y guide target region with a stuffer fragment that could be released by BaeI (NEB) and allow any guide to be inserted easily (p426-crRNA-BaeI). We then used p426-crRNA-BaeI as a template to amplify a fragment containing the new cassette with the *SNR52* promoter and the *URA3* gene using oligos trp1-S-ura3 and Cyc-K-SNR52 (Table S5). This fragment was combined with p414-TEF1p-Cas9-CYC1t that had been cut with SnaBI and Acc65I and assembled using Gibson mix to create p416-TEF1p-Cas9-NLS-crRNA-BaeI. The net effect of these manipulations is to (1) combine the guide RNA and Cas9 genes on a single centromeric (low copy) plasmid, (2) create a flexible entry site for any guide sequence, and (3) replace the *TRP1* marker with *URA3*. To target the *S. cerevisiae YTA12* 5’UTR, we cleaved p416-TEF1p-Cas9-NLS-crRNA-BaeI with BaeI, annealed the YTA12_top YTA12_bot 25-mers together (Table S5) and ligated them to the BaeI cleaved plasmid to produce p416-TEF1-Cas9-NLS-CYC1t-crRNA-YTA12. The advantage of this single plasmid system is that guide sequences are more easily inserted, only one plasmid is needed, and cells lacking the plasmid can be selected after editing on 5-fluororotic acid (5-FOA) plates.

Rescue fragments were created by annealing combinations of synthetic oligonucleotides (Table S5) and filling them in with DNA polymerase. These fragments contained the sequences needed to edit the *S. cerevisiae YTA12* 5’ UTR so that it contained the *S. bayanus* intron, or was deleted of the intron-syntenic sequences, or contained different 5’ ss mutations of the *S. bayanus* intron. Candidate edited yeast clones were grown, and DNA was isolated and analyzed by PCR using primers on either side of the edited site. PCR products were purified and sequenced at the U. C. Berkeley sequencing center to confirm correct editing. Yeast strains determined to contain the correct sequence were streaked on 5-FOA to select clones that have lost the p416-TEF1-Cas9-NLS-CYC1t-crRNA-YTA12 plasmid.

## Acknowledgements

Thanks very much to Joshua Arribere and Russ Corbett-Detig for insightful critical analysis of an earlier draft of this work. We appreciate Sam Fagg, Jen Quick-Cleveland, Stephanie Nystrom, and Santiago Sanchez for their constructive comments on this work. Thanks to Kevin Karplus for suggesting that codon-permutation could be used to create randomized ORF sets that preserve codon usage and GC content. We thank UCLA colleagues Tracy Johnson and Stephen Douglass for discussions and sharing results prior to publication. Thank you to Shana McDevitt and the Vincent J. Coates Genomics Sequencing Laboratory for advice on sequencing library preparation and for sequencing services.

---

**Supplemental Figure 1.**
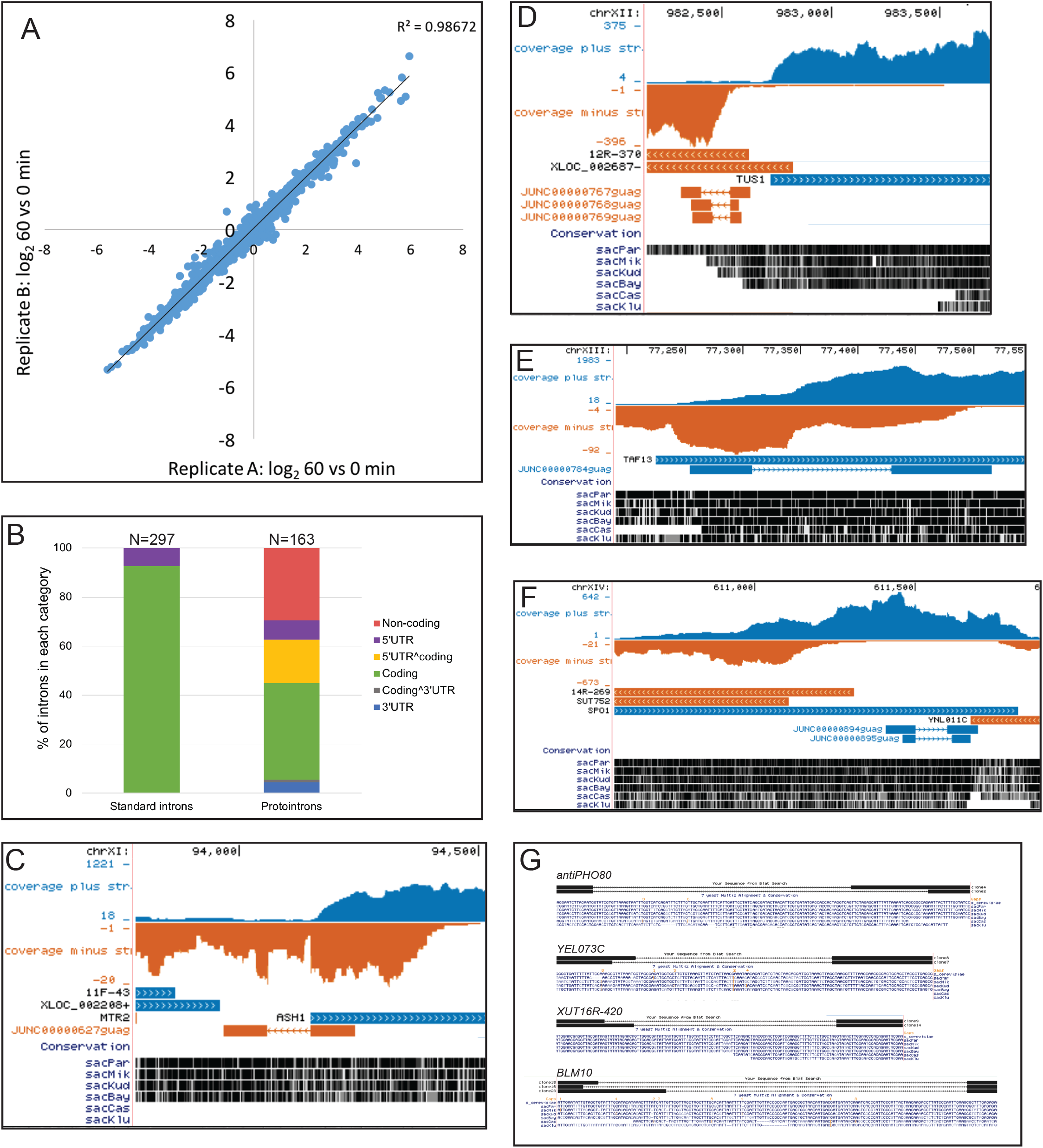
**(A)** Coherence of gene expression changes after 60 minute rapamycin treatment between the two replicate experiments. Log2ratio of treatment to control read coverage over genes was plotted giving an *R*^2^ value of ~0.99. Supplemental to Fig 1A. **(B)** Percentage of standard introns and protointrons that are located in non-coding, 5’UTR, 5’UTR^coding, coding, coding^3’UTR and 3’UTR regions. **(C)** Coverage tracks showing transcription through the genomic locus upstream of *ASH1* where the *antiASH1* protointron is located. Supplemental to Figs 1 B and C. **(D)** Coverage tracks showing transcription through the genomic locus upstream of *TUS1* where the *XUT12R-370* protointron is located. Supplemental to Figs 1 B and C. **(E)** Coverage tracks showing transcription through the genomic locus of *TAF13* where the *TAF13* protointron is located. Supplemental to Figs 1 B and C. **(F)** Coverage tracks showing transcription through the genomic locus of *SPO1* where the *ncSPO1* protointron is located. Supplemental to Figs 1 B and C. **(G)** Alignment of sequenced RT-PCR products showing the location of protointrons with unusual 5’ ss. Supplemental to Fig 1D.

**Table S1:** Saccharomyces cerevisiae standard introns

**Table S2:** *Saccharomyces cerevisiae* overlapping standard introns

**Table S3:** Saccharomyces cerevisiae protointrons

**Table S4:** Saccharomyces cerevisiae filtered reads

**Table S5:** Oligonucleotides

**Table S6:** Saccharomyces mikatae introns

**Table S7:** Saccharomyces bayanus introns

**Table S8:** *Saccharomyces cerevisiae* in-frame protointrons

