## Supplementary figures and images for "Rapidly evolving protointrons in *Saccharomyces* genomes revealed by a hungry spliceosome"

### Supplemental Figure S1

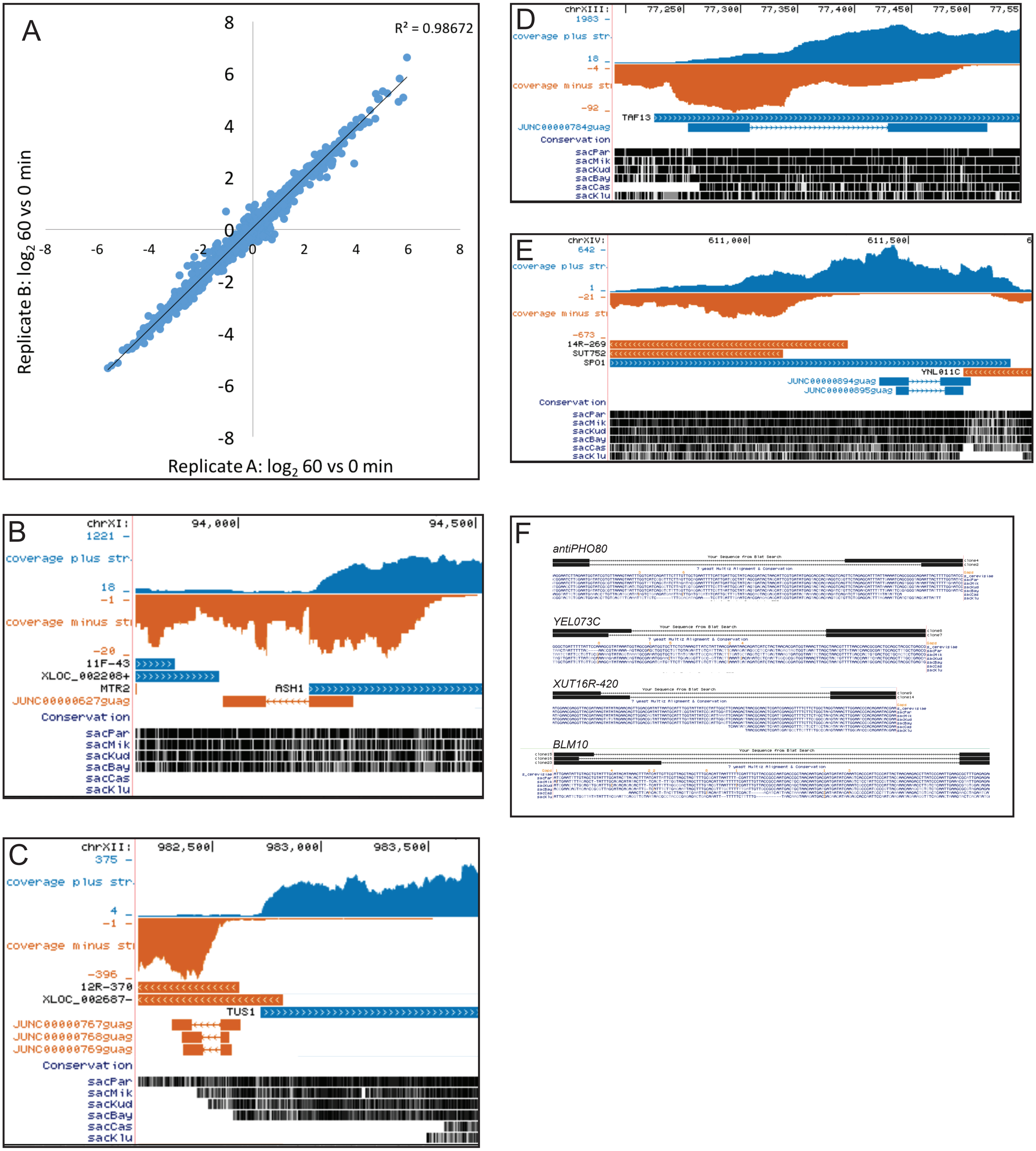
